# A unified developmental framework of the human placenta in its uterine environment *in vivo* and *in vitro*

**DOI:** 10.64898/2026.06.02.729648

**Authors:** Qian Li, Ashley Moffett, Naomi McGovern

## Abstract

Despite advances in single-cell profiling of the human placenta, the genetic programs governing its physiological remodeling throughout gestation remain incompletely understood; this limits the interpretation of trophoblast organoid models. Here, we reconstruct the human placenta in its uterine environment across gestation by integrating public single-cell data into a unified developmental framework developed through a specialized computational strategy. We resolve 100 cell subtypes, expanding the known cellular repertoire and uncovering extensive gestational dynamics. In the placental mesenchymal core, we define a stromal-vascular niche comprising previously unresolved fibroblast heterogeneity and vascular hierarchies (capillary, arterial, and venous). This niche undergoes reprogramming from early angiogenesis to vascular maturation at term and engages signaling programs that support villous homeostasis. Within the trophoblast lineage, we uncover differential progenitor dynamics: bipotent cytotrophoblast (CTB) progenitors persist throughout gestation, whereas extravillous trophoblast (EVT)-biased progenitors are almost absent at term, coinciding with differentiation into specialized states. Benchmarking trophoblast organoids against this reference shows distinct regional identities and developmental biases. Tissue-derived models recapitulate villous CTB whilst trophoblast stem cell-derived organoids resemble smooth chorion CTB; all models capture early gestational syncytiotrophoblast and progressive EVT differentiation. Together, this work provides a resource for understanding placental remodeling across gestation and guiding the use of *in vitro* models.

## Introduction

The placenta is essential to support fetal growth and development with coordinated interactions with both maternal blood in the intervillous space and the uterine mucosa, the decidua^1^. The human placenta has a villous structure with a mesenchymal core populated by fibroblasts, endothelial cells, and distinctive macrophages, known as Hofbauer cells^2^. Surrounding this core is an inner layer of villous cytotrophoblast cells (CTB) and an outer multinucleated syncytiotrophoblast (STB) layer that is covered by maternal blood to mediate nutrient and gas exchange^2^. Extravillous trophoblast cells (EVT) invade the maternal decidua to remodel spiral arteries for establishing uteroplacental blood flow^3^. Away from the definitive placenta, the smooth chorion of the fetal membranes is in contact with the decidua parietalis^4^. The smooth chorion is a layer of trophoblast lacking villi, with no decidual invasion^4^. Beginning in the secretory phase, endometrium is transformed into decidua during pregnancy with differentiation of all cellular elements (stromal, vascular, epithelial, and immune cells) that regulate placentation^5,6^.

This complex cellular ecosystem constitutes a highly dynamic organ, undergoing structural and functional changes throughout gestation^7,8^ (Figure 1A). Early pregnancy relies mainly on histotrophic nutrition derived from decidual gland secretions^9^. There is a crucial transition to haemotrophic nutrition around 8-10 weeks of gestation following the onset of maternal blood flow into the intervillous space^9^. As fetal demands increase, placental growth is accompanied by expansion of the chorionic villous tree with an increased surface area to enhance exchange^7^.

**Figure 1.**
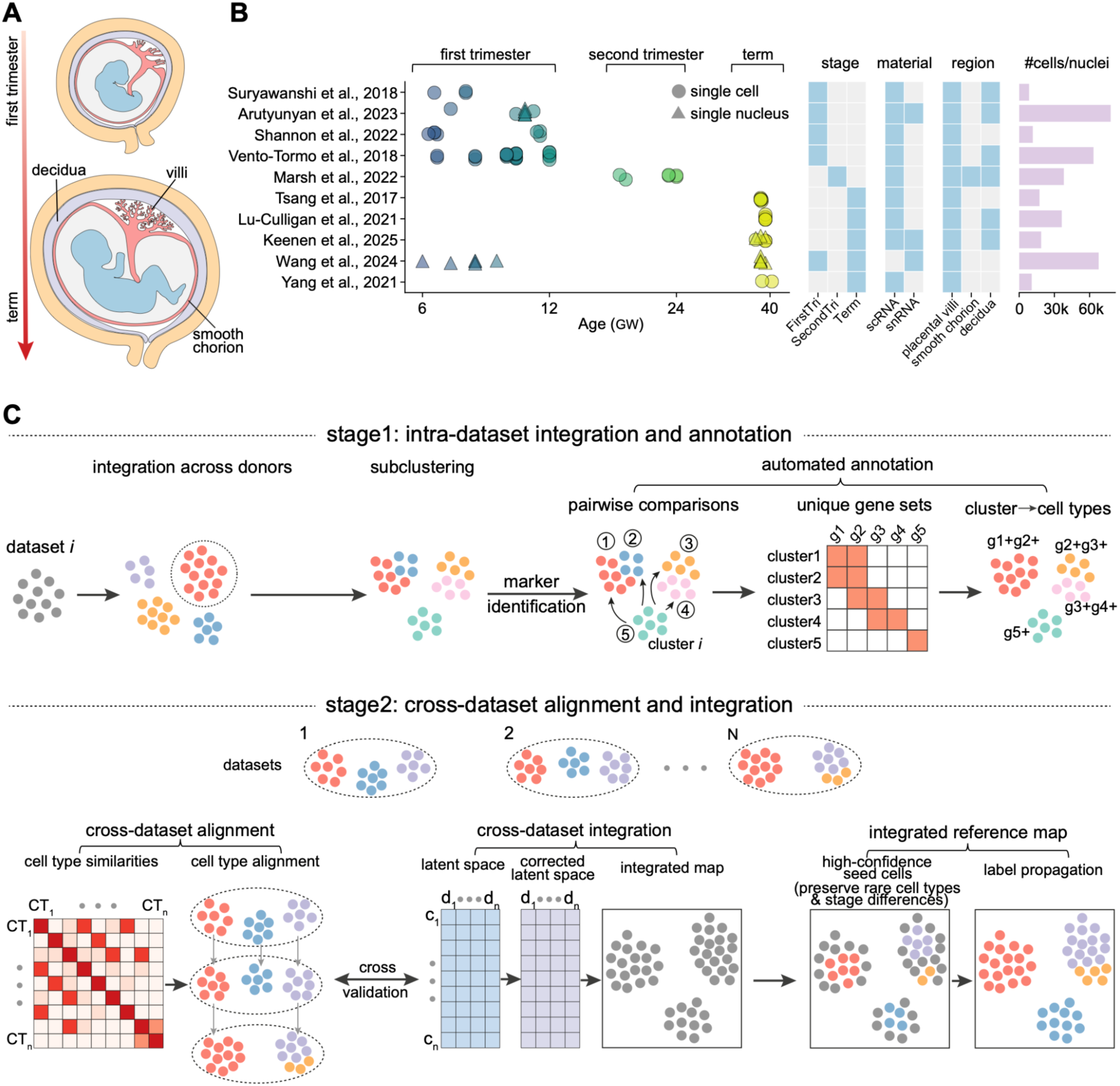
A computational pipeline to preserve developmental dynamics and capture rare cell populations. (**A**) Schematic illustration of human placental development across gestation. (**B**) Summary of publicly available scRNA-seq and snRNA-seq datasets profiling the human placenta and its uterine environment from 10 studies spanning early to late pregnancy. Each dot represents an individual sample, colored by gestational age. (**C**) Overview of the two-stage computational pipeline designed to preserve developmental dynamics and capture rare cell populations. In the first stage, each dataset is analyzed independently to obtain high-resolution cell annotations through subclustering and automated annotation. In the second stage, cross-dataset alignment and integration are performed in parallel to cross-validate cell subtype correspondence across datasets, enabling robust definition of final cell subtypes as well as their high-confidence seed cells for constructing a unified reference map through label propagation.

Despite anatomical knowledge of these processes, the cellular and molecular programs that govern the changes in human placental development across gestation remain elusive. Defining these programs is critical for elucidating both normal placental function and the pathogenesis of pregnancy disorders. Single-cell transcriptomic studies have begun to resolve placental cellular heterogeneity, but most are restricted to specific gestational stages or cell lineages^10–21^. Although a recent study spans early to late pregnancy at the maternal-fetal interface, it lacks systematic characterization of cross-stage dynamics^22^, a task that remains challenging as developmental variation is often confounded with technical batch effects. Consequently, a high-resolution developmental map of the human placenta that captures both cellular diversity and temporal progression is lacking. Addressing this gap requires a specialized computational framework that enables cross-stage integration while retaining stage-specific biological signals.

Such an *in vivo* map is also essential for evaluating trophoblast organoids, which are promising *in vitro* models for experimenting on the human placenta. Trophoblast organoids have been derived from human trophoblast stem cells (TSC-Org)^23^ as well as from first-trimester (FirstTri-Org)^24,25^ and term (Term-Org)^26^ placental tissues. Despite previous efforts to benchmark these models^14,27–29^, the *in vivo* references used are typically restricted to a single developmental stage or anatomical region, thereby limiting comprehensive assessment of their biological relevance.

Here, we assembled publicly available single-cell and single-nucleus RNA sequencing (scRNA-seq and snRNA-seq) datasets spanning multiple gestational stages and placental regions *in vivo*^10,11,13,14,16–19,21,28^, together with three trophoblast organoid models *in vitro*^13,14,27,28^. We complemented these data with spatial transcriptomic profiling using Visium and STARmap in situ sequencing^14,15^. We developed a computational pipeline designed to preserve developmental dynamics and capture rare populations, enabling identification of 100 cell subtypes across gestation. This showed previously underappreciated cellular heterogeneity and developmental dynamics, while providing a framework to systematically evaluate trophoblast organoid models.

## Results

### A computational pipeline to preserve developmental dynamics and capture rare cell populations

To characterize the cellular and molecular changes in the human placenta within its uterine environment across gestation, we collected 10 publicly available scRNA-seq and snRNA-seq datasets from 49 donors covering all stages of gestation (first trimester, second trimester and term) and different regions (placental villi, smooth chorion, and decidua)^10,11,13,14,16–19,21,28^ (Figure 1B and Table S1). Raw data were reprocessed under a unified framework for cell filtering, doublet identification, ambient RNA removal, and maternal or fetal origin inference (Figure S1A), yielding 350,390 high-quality cells for cross-stage analysis.

To address the limitations of existing integration workflows in preserving developmental dynamics and resolving the complexity of the human utero-placental environment, we developed a specialized computational pipeline consisting of two sequential stages: intra-dataset integration and annotation, followed by cross-dataset alignment and integration (Figure 1C).

In the first stage, each dataset was analysed independently to correct donor- and batch-specific effects while maintaining biological variation. Major cell populations were further subclustered to resolve rare populations. We designed an automated annotation workflow to effectively refine subclusters by retaining transcriptionally distinct groups and merging highly similar ones, thereby defining cell subtypes based on unique and robust gene signatures (see Methods; Figure 1C). By analyzing datasets independently, critical differences across developmental stages were retained and incorporated as prior information to guide subsequent cross-dataset integration.

In the second stage, cell types and subtypes were quantitatively aligned across datasets, which enables preservation of cellular heterogeneity across developmental stages. Cross-dataset alignment was further validated through a batch-corrected low-dimensional latent space. High-confidence seed cells for all cell populations present were then defined. Through further label propagation in the latent space, a complete cellular landscape of the human placenta within its uterine environment across gestation was generated (see Methods; Figures 1C and 2A).

**Figure 2.**
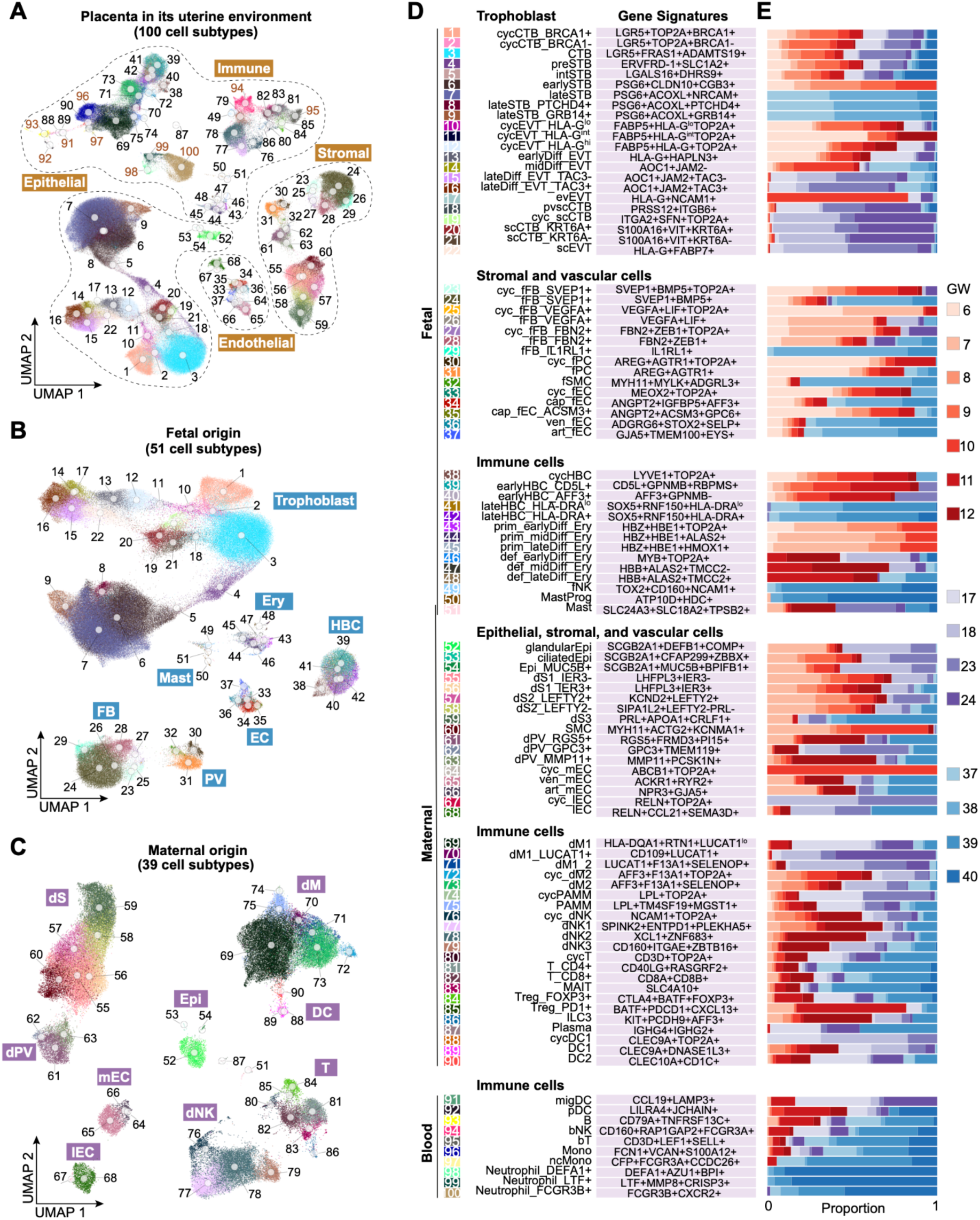
A high-resolution developmental map of the human placenta in its uterine environment throughout gestation. (**A-C**) UMAP visualization of cell subtypes identified across gestation in the human placenta and its uterine environment (**A**), with fetal-derived (**B**) and maternal-derived (**C**) populations also shown separately. Brown numbers in (**A**) indicate blood cell subtypes. FB, fibroblasts; PC, pericytes; SMC, smooth muscle cells; EC, endothelial cells; HBC, Hofbauer cells; Ery, erythrocytes; NK, natural killer cells; Epi, epithelial cells; S, stromal cells; PV, perivascular cells; M, macrophages; PAMM, placenta-associated maternal macrophages; DC, dendritic cells; Mono, monocytes; cyc, cycling; lo, low; int, intermediate; hi, high; Diff, differentiation; ev, endovascular; sc, smooth chorion; pvsc, placental villi and smooth chorion; f, fetal; cap, capillary; ven, venous; art, arterial; prim, primitive; def, definitive; Prog, progenitor; d, decidual; m, maternal; l, lymphatic; b, blood; nc, non-classical. (**D**) Harmonized nomenclature and molecular signatures of each cell subtype, with numbers shown on the left corresponding to those in (**A-C**). (**E**) Relative abundance of cells from different developmental stages in each cell subtype, with distributions across individual gestational age points within each stage also indicated.

### A high-resolution developmental map of the human placenta in its uterine environment throughout gestation

Through the specialised computational pipeline, we identify 100 subtypes in the human placenta and its uterine environment, substantially expanding the known cellular repertoire of this niche (Figure 2A). This developmental map covers all three trimesters (Figure S1B) and spans the epithelial, stromal, endothelial and immune compartments (Figure 2A). Amongst these, 51 subtypes are of fetal origin (Figures 2B and S1C), 39 of maternal origin (Figures 2C and S1C), and 10 are probably circulating blood cell types (Figures 2A and S1C). We have provided harmonized cell labels within the developmental context to resolve discrepancies across studies, with each subtype being further molecularly characterized by a unique combination of gene signatures (Figures 2D and S2A).

In the placenta, we identify 22 trophoblast subtypes, 15 stromal and vascular subtypes, and 14 immune subtypes of fetal origin (Figures 2D and S2A), revealing extensive cellular remodeling across gestation (Figure 2E). For example, Hofbauer cells (HBC) undergo substantial temporal changes: cycling HBC, *CD5L*⁺ HBC, and *AFF3*⁺ HBC are enriched in the first trimester, whereas *HLA-DR*⁺ HBC predominate at term (Figures 2E and S2B), consistent with our previous findings^30^. In addition, we detect both primitive and definitive erythrocytes in the placenta (Figures 2D and S2A), with primitive erythrocytes largely present before 10 gestational weeks and definitive erythrocytes emerging thereafter (Figure 2E). This observation firmly establishes the placenta as an additional site of primitive hematopoiesis^31,32^. Additionally, several rare placental populations are present, including mast cells and fetal natural killer cells (fNK) that appear at term (Figures 2D, 2E, S2A and S2B).

In the uterine environment, we find 17 epithelial, stromal, and endothelial subtypes and 22 immune subtypes; these all also show dynamic changes throughout gestation. For example, the key decidual immune cells, decidual NK cells (dNK), known to play a role in regulating placentation early in pregnancy^29,33^, predominate in the first trimester and decline thereafter (Figures 2E and S2B). Unlike the other dNK subsets (dNK1 and dNK2), dNK3 are present throughout gestation (Figures 2E and S2B), confirming they are the dominant dNK population at term^34^. In contrast, T cells represent a minor population during early gestation (Figure S2C), with their relative abundance increasing at term, coinciding with the decline in dNK populations (Figures 2E, S2B, and S2C). Within the T cell compartment, two previously described regulatory T cell (Treg) subtypes are found in the decidua^35^: *FOXP3*⁺ and *PD1*⁺ (Figures 2D and S2A). *PD1*⁺ Tregs are enriched in the first trimester, suggesting an important mechanism for T cell tolerance to invading EVT which expresses *PD-L1* (Figures 2E and S2D). Tolerance is also reinforced by DC1, which are more abundant in the first two trimesters and express Gelsolin (*GSN*) and *ETV3* (Figures 2E, S2B and S2D), key factors implicated in dampening T cell responses^36,37^.

Given the extensive anatomical and functional changes occurring in the placenta, we next focus on this environment to further dissect its underlying cellular and molecular organization throughout gestation.

### Dynamic changes in placental stromal and vascular cells that support villous homeostasis

The villous mesenchymal core contains diverse stromal and vascular cell populations that remain poorly characterized. Here, we systematically defined the cellular composition and temporal dynamics of this mesenchymal compartment throughout gestation, revealing considerable heterogeneity and stage-specific specialization.

We identify four non-cycling, transcriptionally distinct fetal fibroblast (fFB) subtypes with dynamic representation across pregnancy (Figures 3A-3C, S3A and S3B). The most abundant population (*SVEP1*^+^) is present throughout gestation (Figures 2E and S3B). In contrast, two subtypes are preferentially enriched in the first trimester. One (*VEGFA*^+^) is characterized by the expression of *VEGFA* and *LIF*, whereas the other (*FBN2*^+^) is enriched for genes associated with extracellular matrix (ECM) (Figures S2A, S3B and S3C). Thus, these populations probably contribute to vasculogenesis and establishment of the villous ECM architecture in early pregnancy (Figure S3C), consistent with upregulation of angiogenic and ECM programs in the first trimester within fFB cells (Figure 3D). The three subtypes described above each have their own proliferating pools in the first trimester (Figures 3B and S3B), further indicating their distinct origins and identities. A fourth subtype (*IL1RL1*^+^) emerges exclusively at term (Figures 2E and S3B). IL1RL1 (ST2) is the receptor for IL-33 that is abundantly expressed by endothelial cells and released upon cellular stress or damage^38–41^ (Figure S3D). Additionally, at term, all fFB subtypes upregulate genes associated with antioxidant defence (Figure 3D), offering potential protection against oxidative stress.

**Figure 3.**
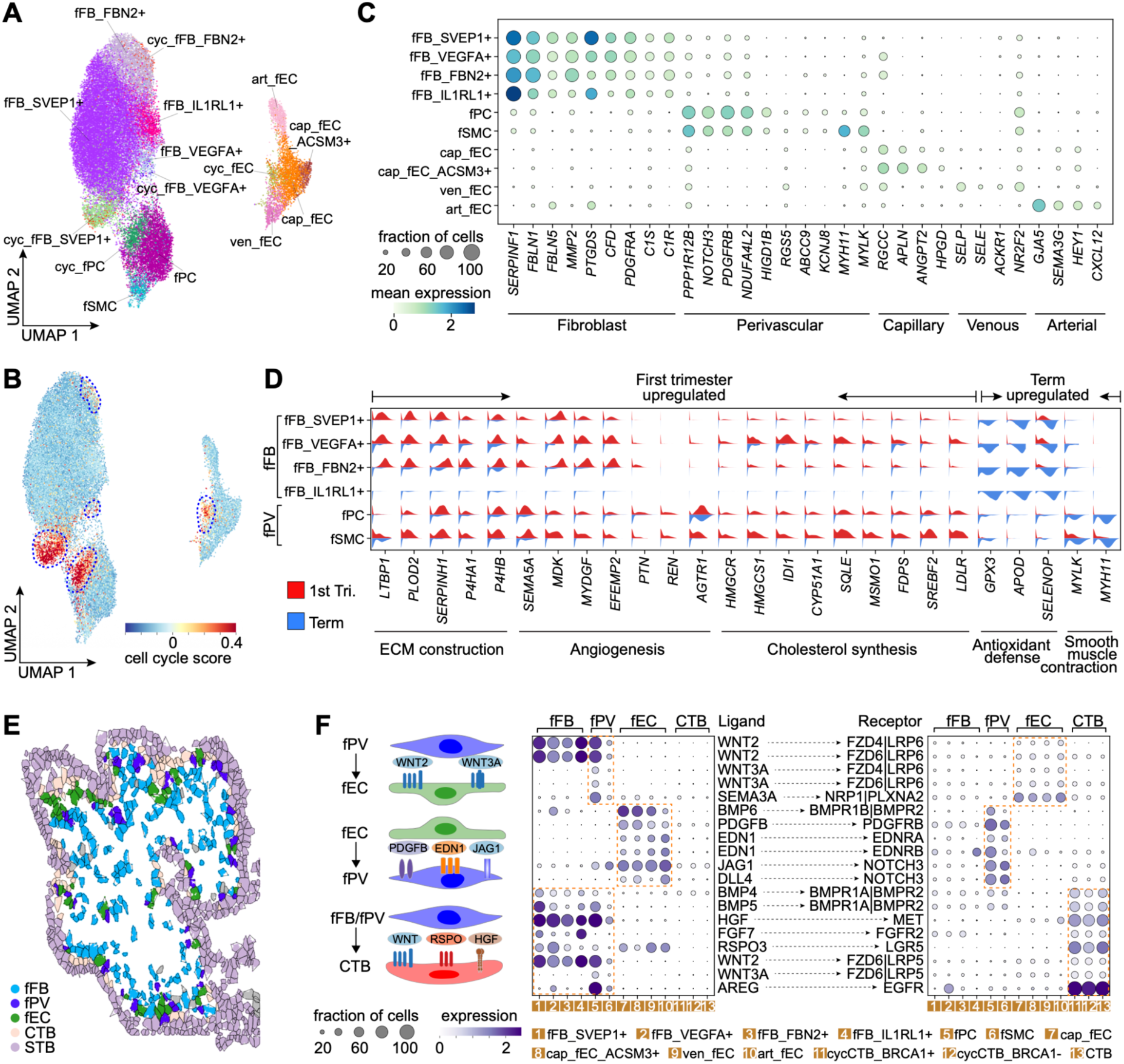
Dynamic changes in placental stromal and vascular cells that support villous homeostasis. (**A**) UMAP visualization of stromal and vascular subtypes identified across gestation within the mesenchymal core of the placental villi. (**B**) UMAP visualization of cell cycle scores calculated based on the expression of cell cycle-associated genes. Proliferative populations are highlighted by dotted blue circles. (**C**) Dot plot showing expression of marker genes for fibroblast, perivascular, capillary, venous, and arterial endothelial cells across placental stromal and vascular subtypes. Dot color and size indicate average expression level and proportion of cells expressing the genes, respectively. (**D**) Violin plots showing expression of genes associated with distinct biological processes and enriched in either first-trimester or term fibroblast and perivascular cells. Expression levels in cells from the first trimester (red) and term (blue) are shown for each fibroblast and perivascular subtype. PV, perivascular cells. (**E**) Spatial distribution of placental cell types within a placental villus predicted using the developmental reference map. (**F**) Ligand-receptor-mediated interactions between stromal, vascular, and trophoblast populations in the placenta, with the expression of the ligands and receptors across cell subtypes shown separately on the left and right. Dot color and size indicate average expression level and proportion of cells expressing the genes, respectively. A schematic summary of the predicted interactions is shown on the far left.

The placenta receives deoxygenated fetal blood via the two umbilical arteries and oxygenated blood returns to the fetus by the umbilical vein^42,43^. The cells of the villous circulatory system have not been extensively characterized at single-cell resolution. We identify cycling, capillary (two subtypes), arterial, and venous fetal endothelial cells (fEC) throughout gestation (Figures 3A-3C and S3A). Cycling fEC are enriched in the first trimester, consistent with active angiogenesis during early placental development^44^ (Figures 2E and S3B). One capillary subtype, marked by *ACSM3* expression, is also preferentially enriched in the first trimester (Figures 2E and S3B). In contrast, the other capillary, arterial, and venous fEC increase in abundance toward term, reflecting expansion and maturation of the vascular network^44^ (Figures 2E and S3B). The placental circulation is unusual in lacking neural regulation^45^. The pericytes surround capillaries, and smooth muscle cells (SMC) encircle the larger stem vessels^44^. In parallel with endothelial diversification, we identify previously uncharacterized perivascular populations that align with this vascular hierarchy: pericytes associated with capillaries, and SMC surrounding arteries and veins (Figures 3C and S3A). The pericytes identified here have been misclassified as fFB in previous studies^10,11,15^. These perivascular cells, along with fFB, highly express genes encoding key enzymes and transcription factors involved in cholesterol biosynthesis during early gestation (Figure 3D), suggesting that the placenta can synthesize cholesterol de novo in addition to acquiring it from the maternal circulation. They also transition from angiogenic programs in early gestation to smooth muscle contractile programs at term, including upregulation of *MYLK* and *MYH11*, reflecting acquisition of a differentiated SMC phenotype in late gestation (Figure 3D). Dedifferentiation of SMC has been associated with intrauterine fetal growth restriction, highlighting the relevance of vascular maturation programs to placental pathology^46^.

To define spatial relationships within the mesenchymal core, we projected their single-cell transcriptional profiles onto publicly available spatial transcriptomics data generated using STARmap in situ sequencing from early human placentas^15^ (Figure 3E). Perivascular populations localize adjacent to fEC, supporting their anatomical identity (Figure 3E). These stromal and vascular populations are also anatomically close to trophoblast cells, suggesting extensive communication in the villous core (Figure 3E). Ligand-receptor analysis shows reciprocal communication networks within the villi (Figure 3F). fFB and pericytes are predicted to signal to capillary fEC through WNT pathways, for promoting and maintaining angiogenesis^47^, while, capillary fEC recruit pericytes through the PDGFB-PDGFRB signaling axis^48^. Arterial and venous fEC, on the other hand, express *EDN1*, encoding the potent vasoconstrictor peptide^49^ whose cognate receptor, *EDNRA*, is present in SMC. All fEC use the Notch pathway for communicating with perivascular cells, consistent with its role in regulating vascular interactions^50^. Beyond vascular support, fFB and perivascular populations interact with trophoblast cells through secreting growth factors and hormones (Figure 3F). fFB and pericytes are major sources of HGF and WNT ligands, whose receptors are expressed by CTB, suggesting a role in supporting trophoblast proliferation^51^. Distinct fFB subtypes also have specialized and stage-specific secretory programs: *SVEP1*^+^ and *VEGFA*^+^ fFB show elevated R-spondin (*RSPO3*) during early gestation, a crucial factor for maintaining CTB stemness^24,52–54^; *SVEP1*^+^ fFB is also the dominant source of *BMP4* and *BMP5*, particularly at later stages, which are regulators of trophoblast development and differentiation^55–57^; and the term-specific *IL1RL1*^+^ fFB expresses *FGF7*, a factor also expressed by other fFB subtypes specifically at term, potentially promoting CTB survival in late gestation^58–60^ (Figures 3F and S3E). Collectively, these findings reveal dynamic reprogramming of the stromal and vascular cells that supports vascular maturation and trophoblast homeostasis throughout gestation.

### Sequential emergence of distinct trophoblast subsets and differential persistence of their progenitors throughout gestation

Trophoblast cells constitute a major cellular compartment of the placenta, yet their full heterogeneity and developmental dynamics across gestation remain incompletely defined. Using our gestationally resolved reference map, we identify 22 trophoblast subtypes: three CTB, six STB, and eight EVT subtypes, as well as five subtypes from the smooth chorion (Figures 4A and S4A), providing a higher-resolution view accompanied by a harmonized, developmentally-contextualized nomenclature (Figures 2D and 4A). Integration of snRNA-seq data enables reconstruction of the full STB differentiation continuum from upstream CTB through pro-fusing intermediates to mature STB (Figure 4A), as well as the dissection of gestational changes in mature STB. We identify four mature STB subtypes, including one subtype (*CGB3*+) specific to early gestation and three enriched at term (*ACOXL*+*PTCHD4*–*GRB14*–, *PTCHD4*+, *GRB14*+) (Figures 2D, 2E, S2A and S4B). The term *GRB14*+ subtype also expresses *GPC5*, known to be enriched in syncytial knots^22^. First-trimester STB are transcriptionally distinct from term ones (Figure S4C), reflecting previously reported stage-specific specialization^21^. For example, *CGB3*, a subunit of human chorionic gonadotropin (hCG), is enriched in the first trimester and absent at term (Figure S4C), consistent with hCG secretion dynamics across gestation and previous reports^28,61^. In contrast, *MFSD2A*, both a transporter of docosahexaenoic acid (DHA) needed for fetal brain development and a receptor for syncytin-2 required for STB formation^62–65^, is more highly expressed at term (Figure S4C). We detect multiple cycling trophoblast populations: two CTB, three EVT, and one smooth chorion CTB cycling subtypes (Figures 4A and 4B). These populations show graded proliferative capacity and distinct lineage biases. *BRCA1*^+^ cycling CTB has the highest proliferative signature and is positioned at the root of the entire trophoblast differentiation trajectory by both pseudotime and RNA velocity analyses (Figures 4B, 4C and S4D). Located downstream of *BRCA1*^+^ cycling CTB, *BRCA1*^−^ cycling CTB has reduced proliferative activity, suggesting a more lineage-restricted state (Figures 4B, 4C and S4D). The cycling CTB found in the smooth chorion strongly express *ITGA2* (Figure S2A), a marker previously associated with a small subset of cells in the cytotrophoblast columns^66^. The three cycling EVT populations have different levels of *HLA-G* expression, from barely detectable to robust expression (Figure S2A). This correlates with progressively diminished proliferation, stepwise loss of CTB and gradual acquisition of EVT gene signatures (Figure 4B), consistent with progressive commitment toward EVT identity (Figures 4C and S4D). Trajectory inference suggests a refined trophoblast differentiation model in which cycling CTB generate non-cycling CTB which in turn give rise to STB. In contrast, cycling CTB act as upstream progenitors that generate cycling EVT and subsequently EVT (Figures 4C and S4D). This is supported by the close spatial proximity of cycling CTB and cycling EVT within the proximal cell column near the villous core in the spatial mapping (Figure 4D). Together, these data resolve trophoblast progenitor organization into a structured continuum rather than discrete, isolated states.

**Figure 4.**
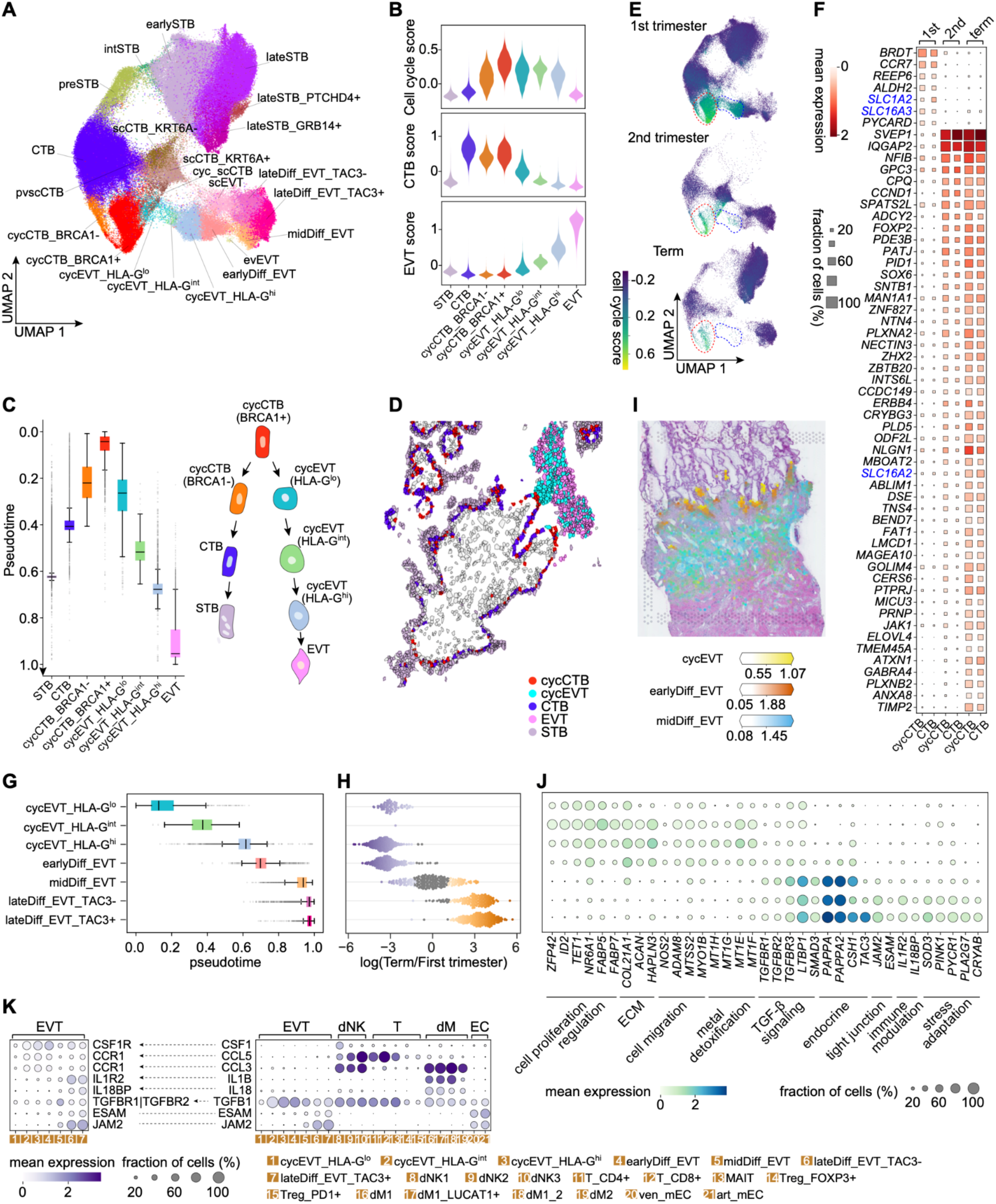
Sequential emergence of distinct trophoblast subsets and differential persistence of their progenitors throughout gestation. (**A**) UMAP visualization of trophoblast subtypes identified across gestation in the placental villi and smooth chorion. (**B**) Violin plots showing distributions of cell cycle, CTB, and EVT scores across trophoblast populations. Cell cycle scores were calculated based on expression of cell cycle-associated genes, whereas CTB and EVT scores were derived from corresponding lineage-specific gene signatures. (**C**) Box plot showing inferred pseudotime across trophoblast populations, with the proposed trophoblast differentiation model shown on the right. Center lines, hinges, and whiskers indicate medians, interquartile ranges, and 5th, 95th percentiles, respectively. (**D**) Spatial distribution of trophoblast subtypes within an anchoring villus predicted using the developmental reference map. (**E**) UMAP visualization of cell cycle scores in trophoblast cells from the first trimester (top), second trimester (middle), and term (bottom). Cycling CTB and cycling EVT populations are highlighted by red and blue dotted circles, respectively. (**F**) Expression of first-trimester-enriched and top 50 term-enriched protein-coding genes in cycling and non-cycling CTB cells across gestational stages. Transporter genes are highlighted in blue. Square color and size indicate average expression level and proportion of cells expressing the genes, respectively. (**G**) Box plot showing inferred pseudotime across EVT subtypes. Center lines, hinges, and whiskers indicate medians, interquartile ranges, and 5th, 95th percentiles, respectively. (**H**) Changes in abundance of EVT subtypes between the first trimester and term, with y-axis labels shared with (**G**). Each dot represents a neighborhood assigned to a subtype, and colored dots indicate significant abundance changes (FDR < 0.1). (**I**) Spatial distribution of EVT subtypes at the maternal-fetal interface predicted using the developmental reference map. Color gradient indicates cell densities inferred by cell2location. (**J**) Dot plot showing expression of genes associated with distinct programs and enriched in EVT subtypes at different stages of differentiation, with y-axis labels shared with (**G**). Dot color and size indicate average expression level and proportion of cells expressing the genes, respectively. (**K**) Predicted ligand-receptor-mediated interactions between EVT and decidual populations, with the expression of the ligands and receptors across cell subtypes shown separately on the right and left. Dot color and size indicate average expression level and proportion of cells expressing the genes, respectively.

Cycling trophoblast populations have distinct gestational dynamics. While cycling CTB decline in abundance over time (Figures 2E and S4B), they are still present at term (Figure 4E), indicating sustained maintenance of CTB progenitors throughout pregnancy. The downstream non-cycling CTB are similarly preserved across gestation, although sparser at term (Figures 4E and S4B). They also undergo molecular changes over time: differential expression analysis identifies 7 and 153 protein-coding genes enriched in the first trimester and at term, respectively, indicating more extensive transcriptional remodeling at term (Figure 4F). Among these, transporter genes show dynamic expression across gestation. For example, *SLC1A2* and *SLC16A3*, involved in glutamate and lactate transport^67–69^, have higher expression in the first trimester. At term, *SLC16A2* (MCT8), a specific transporter for thyroid hormone, is upregulated, consistent with its role in mediating transplacental transfer of thyroid hormones to support fetal development^70^.

In contrast to cycling CTB, cycling EVT progressively diminish and are almost absent at term, suggesting stage-restricted progenitor activity for EVT (Figure 4E). Further integration of differential abundance analysis, trajectory inference, and spatial projection using publicly available Visium data^14^ shows both temporal specification and spatial stratification of EVT (Figures 4G-4I). EVT at earlier stages of differentiation are enriched in the first trimester and localized proximally in the cell column adjacent to villi, whereas more differentiated EVT predominate at later gestational stages and are positioned within the decidua. These findings are in keeping with early gestation as the major window for EVT expansion and decidual invasion associated with spiral artery remodeling^71^.

Comparative molecular analysis further reveals stage-specific gene programs in EVT (Figures 4J and 4K). EVT at earlier stages of differentiation, which are enriched in the first trimester (hereafter referred to as early EVT), show gene programs associated with regulation of cell proliferation, ECM organization, and cell migration, consistent with their proliferative and invasive phenotype (Figure 4J). These early EVT also upregulate metallothioneins (*MT1H, MT1G, MT1E,* and *MT1F*), which bind heavy metals such as cadmium and zinc, providing potential protection against environmental toxins and supporting proper differentiation^72^. EVT differentiation is also supported by elevated expression of cytokine receptors in early EVT (Figure 4K), indicative of responsiveness to dNK-derived cytokines known to promote EVT differentiation^29^ (Figure 4K). In contrast, EVT at late stages of differentiation, which are enriched towards term (hereafter referred to as late EVT), show gene programs consistent with a differentiated state. For example, they are enriched for TGF-β signaling components^73^, including *TGFBR1* and *TGFBR2* (Figure 4J), predicted to bind to TGF-β ligand produced by EVT, decidual immune and endothelial cells (Figure 4K). They also show gene signatures associated with growth factor regulation (*PAPPA*, *PAPPA2*) and endocrine factors (*CSH1*) (Figure 4J). *TAC3*, specifically enriched in late EVT, encodes a vasoactive neuropeptide^74^, pointing to a role in regulating uterine blood flow later in gestation. In parallel, they upregulate endothelial-associated genes involved in tight junction pathways (Figures 4J and 4K). Late EVT also express decoy receptors, including *IL1R2* and *IL18BP*^75^ (Figure 4J), which neutralize pro-inflammatory IL-1β and IL-18 produced by decidual macrophages (Figure 4K). Moreover, oxidative stress-adaptive genes are observed in late EVT (Figure 4J). Collectively, EVT show substantial remodeling across gestation, demonstrating diverse roles such as endocrine, vascular, immune-modulatory, and stress-adaptive functions, particularly later in pregnancy.

### Trophoblast organoids differentially recapitulate regional and developmental trophoblast states *in vivo*

The trophoblast diversity we have captured *in vivo* across gestation and placental regions (villous placenta and smooth chorion) has allowed us to systematically benchmark the different trophoblast organoid models. We collected four publicly available scRNA-seq and snRNA-seq datasets generated from tissue-derived organoids (FirstTri-Org and Term-Org) and TSC-derived organoids (TSC-Org) across 18 donors^13,14,27,28^ (Figure 5A). Raw data were reprocessed using the same unified framework applied to primary tissue (Figure S1A), followed by integration across datasets and additional quality control. In total, 131,421 cells and nuclei are retained and classified into 13 trophoblast subtypes corresponding to CTB, STB, EVT, and smooth chorion CTB, based on gene signatures *in vivo* and label transfer from the primary trophoblast reference map (Figures 5B-5E and S5A-S5C).

**Figure 5.**
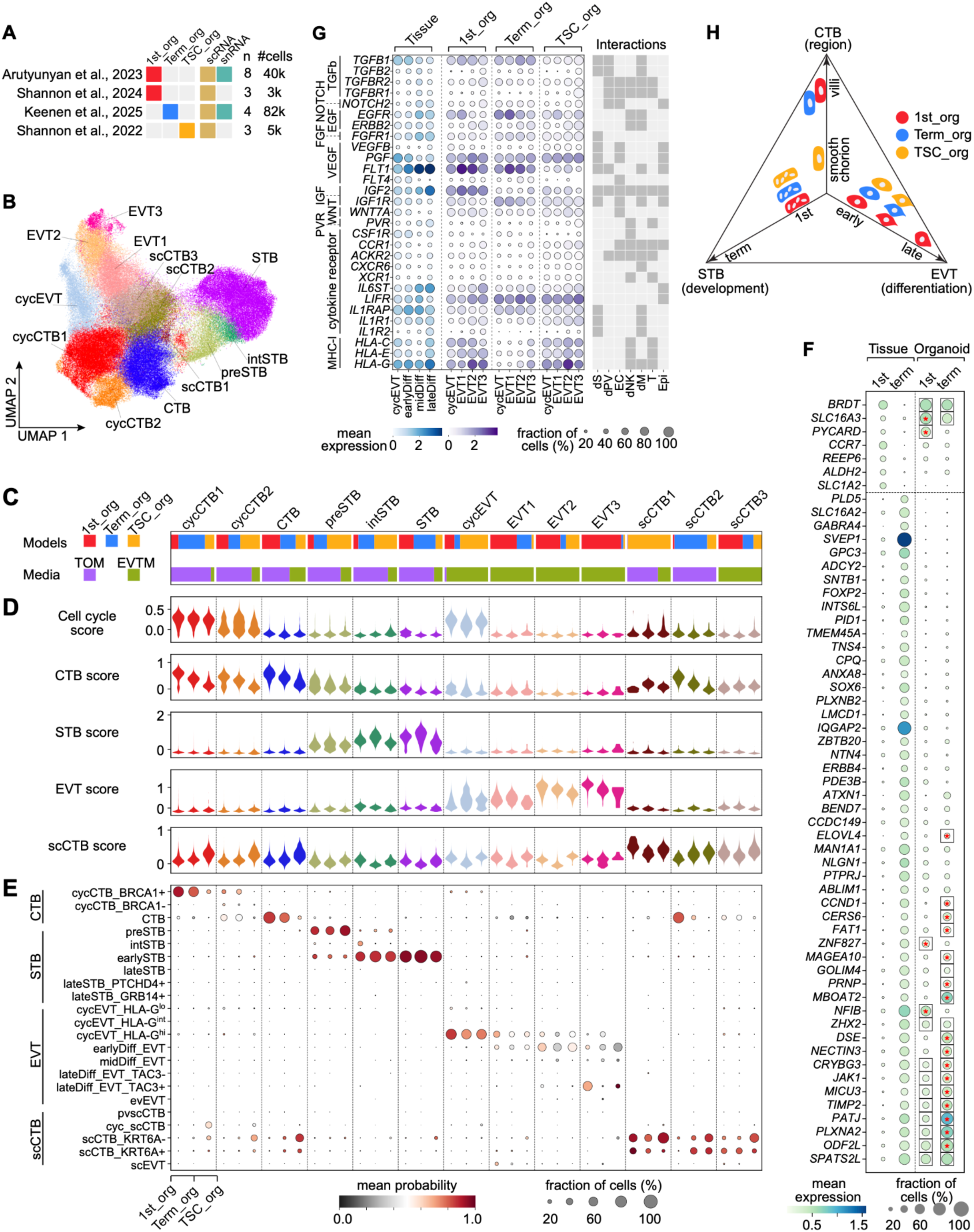
Trophoblast organoids differentially recapitulate regional and developmental trophoblast states *in vivo*. (**A**) Summary of publicly available scRNA-seq and snRNA-seq datasets generated from the three trophoblast organoid models. *n*, number of donors. (**B**) UMAP visualization of trophoblast subtypes identified across trophoblast organoids. (**C**) Distribution of cells from different organoid models (top) and culture media (bottom) in each trophoblast subtype. TOM, trophoblast organoid medium; EVTM, EVT differentiation medium. (**D**) Violin plots showing distributions of cell cycle, CTB, STB, EVT, and scCTB scores across different organoid models in each trophoblast subtype. Cell cycle scores were calculated based on expression of cell cycle-associated genes, whereas lineage scores were derived from the corresponding top 50 *in vivo* gene signatures. (**E**) Predicted cell identities for *in vitro* cell subtypes from each organoid model based on the *in vivo* developmental reference map. Dot color and size indicate prediction probability and proportion of *in vitro* cells aligned to each *in vivo* subtype, respectively. (**F**) Dot plot showing expression of first-trimester- and term-enriched genes identified from *in vivo* CTB across the primary tissue and tissue-derived organoids. Squares indicate expression in >30% of cells in the organoids, and asterisks indicate significant expression differences between FirstTri-Org and Term-Org. Dot color and size indicate average expression level and proportion of cells expressing the genes, respectively. (**G**) Dot plot showing expression of key ligand-receptor components of major pathways involved in EVT-decidua interactions across EVT subtypes *in vivo* and *in vitro*. The heatmap on the right shows the corresponding decidual populations predicted to interact with EVT through each signaling component. Dot color and size indicate average expression level and proportion of cells expressing the genes, respectively. (**H**) A schematic summary of *in vivo* trophoblast states recapitulated by the three trophoblast organoid models.

We identify two cycling and one non-cycling CTB subtypes enriched for *in vivo* CTB signatures in all these models (Figures 5B-5D). The two cycling CTB populations show graded proliferative capacity, mirroring the heterogeneity observed *in vivo* (Figure 5D). HLA class I molecules, always absent in *in vivo* CTB, have model-specific patterns. Tissue-derived organoids (FirstTri-Org and Term-Org) show minimal expression of *HLA-A* and *HLA-B*, whereas we confirm that TSC-Org display detectable *HLA-B* expression^76^ (Figure S5B). We also find distinct regional identities in the *in vitro* CTB populations across models. CTB from tissue-derived organoids (FirstTri-Org and Term-Org) map predominantly to proliferative and non-proliferative villous CTB *in vivo* (Figures 5E and S5C). In contrast, CTB from TSC-derived organoids show strong smooth chorion CTB (scCTB) signatures (Figure 5D) and align transcriptionally with proliferative and non-proliferative scCTB populations *in vivo* (Figures 5E and S5C). An scCTB-like subtype (scCTB1) corresponding to *in vivo* scCTB is also enriched in TSC-Org (Figures 5C-5E and S5C). Two additional scCTB-like subtypes are also detected in tissue-derived organoids (Figures 5C-5E and S5C). scCTB2 predominates in Term-Org, whereas scCTB3 is present in all three models cultured in EVT differentiation medium (EVTM), suggesting an intermediate state induced during EVT specification.

We next compared CTB from FirstTri-Org and Term-Org as they both recapitulate villous trophoblast states. CTB from FirstTri-Org show stronger *in vivo* villous CTB signatures than Term-Org (Figure 5D), indicating better preservation of primary trophoblast identity. We further assessed the developmental states captured by these models by examining expression of first-trimester- and term-enriched genes identified from *in vivo* villous CTB within these organoids (Figure 5F). Among the top 50 term-enriched genes, 19 (38%) are detected in Term-Org, compared with 11 (22%) in FirstTri-Org (Figure 5F). Thus, Term-Org show some partial acquisition of term-associated transcriptional programs, which are comparatively underrepresented in FirstTri-Org.

Three STB subtypes with progressively increasing STB signatures are detected in all organoid models, corresponding to *in vivo* states from pro-fusion intermediates to mature STB (Figures 5C-5E and S5C). The STB in organoids resemble first-trimester rather than term STB (Figures 5E and S5D), indicating that organoids preferentially support an early gestational STB phenotype.

Four EVT subtypes are present in all organoid models cultured in EVTM, including one cycling population and three subtypes with progressively increasing EVT gene programs, aligned with the EVT differentiation pathway *in vivo* (Figures 5C-5E and S5C). One subtype (EVT3) maps to *in vivo* EVT at a late stage of differentiation and expresses marker genes such as *TAC3* and *JAM2* (Figures 5E, S5B and S5C). EVT3 is enriched in FirstTri-Org (Figure 5C), possibly reflecting prolonged culture in differentiation medium in this model^14^ or intrinsic differences in differentiation capacity. Given the central role of EVT in mediating interactions with decidual cells, we further assessed whether key ligand-receptor components involved in EVT-decidua communication are maintained in organoids. All three models preserve signaling elements of major *in vivo* pathways, including those involved in immune and vascular interactions (Figure 5G), supporting their potential for investigating cellular communication between the placenta and its uterine environment.

To summarize, benchmarking the tissue-derived and TSC-derived trophoblast organoids against our gestationally and regionally resolved *in vivo* reference shows that these models have distinct regional identities and developmental biases (Figure 5H): tissue-derived models recapitulate villous CTB and TSC-derived organoids resemble smooth chorion CTB, while all models preferentially capture early gestational STB and support progressive EVT differentiation.

## Discussion

A successful human pregnancy depends on the coordinated development of the placenta and the establishment of tightly regulated communication networks with its uterine environment. This ecosystem is both structurally complex and temporally dynamic, adapting continuously to the evolving physiological demands of the fetus throughout gestation. Despite previous efforts to dissect the human placenta at single-cell resolution, the genetic programs underlying its dynamic remodelling remain poorly understood. Here, by integrating publicly available single-cell and single-nucleus transcriptomic data spanning multiple gestational stages and anatomical regions, complemented with spatial mapping, we establish a gestationally and regionally resolved reference of the human placenta in its uterine environment at improved resolution. This provides a systems-level view of cellular and molecular remodeling across gestation and offers a unified framework for evaluating *in vitro* models of human placentation within developmental and anatomical contexts.

An important contribution of this work is the development of a computational strategy to address a major challenge in reconstructing developmental systems from heterogeneous public datasets. In highly dynamic organs such as the placenta, the developmental stage itself represents a dominant source of biological variation, making conventional integration approaches prone to overcorrection and loss of biologically meaningful differences^77^. Our two-stage strategy overcomes this limitation by combining dataset-level annotation with cross-dataset alignment, thereby prioritizing preservation of biological variation both within and across datasets. This approach retains stage-specific signals and resolves rare populations that are frequently obscured during global integration, thus providing a generalizable strategy for investigating other dynamic biological systems apart from the placenta and its uterine environment.

Applying this computational strategy, we have constructed a high-resolution developmental map of the human placenta within its uterine environment, comprising 100 subtypes with harmonized nomenclature defined in a developmental context. This is a more diverse cellular ecosystem than previously appreciated, particularly obvious in the villous mesenchymal core. The stromal and vascular organization mirrors anatomical descriptions of placental villi and includes previously uncharacterized fFB heterogeneity together with a vascular hierarchy of capillary, arterial, and venous fEC and their associated pericytes and SMC. fFB emerge as key regulators of vascular development and function. One first-trimester-enriched fFB subtype expresses *VEGFA*, consistent with a role in angiogenesis^78,79^. A term-specific subtype expresses high levels of *IL1RL1* (ST2), functioning as either a transmembrane or decoy receptor for IL-33^80^, a stress-induced factor released by EC^38–41^. This signaling axis suggests a role for this fFB subtype in sensing endothelial stress and maintaining vascular integrity in late gestation. Dysregulation of *IL1RL1* has also been implicated in pregnancy complications like pre-eclampsia and preterm birth^81–83^. In parallel, fFB establish extensive signaling interactions with trophoblast cells through WNTs, HGF, BMPs, R-spondins, and FGFs. These signaling interactions resemble mesenchymal niches in other organs that support epithelial homeostasis and tissue morphogenesis^84–88^. Our findings suggest that placental villous development shares organizational principles with branching morphogenesis in organs such as the lung, providing a framework for understanding how villous architecture and vascular network are established during pregnancy.

Although trophoblast cells have been the primary focus of previous studies, their developmental dynamics across gestation remain incompletely understood. We identify 22 trophoblast subtypes spanning all three trimesters and different anatomical regions, the villous placenta, decidua and smooth chorion. Our analysis refines current models of trophoblast differentiation by delineating both lineage hierarchy and temporal restriction of progenitor activity. Two cycling CTB populations with graded proliferative capacity occupy the root of differentiation trajectories and, although reduced at term, persist throughout gestation, suggesting sustained progenitor maintenance. This explains the successful derivation of trophoblast organoids from term placental tissues^26^. In contrast, EVT progenitors are largely restricted to early gestation. Although EVT have traditionally been considered important in early gestation^71,89,90^, we show that EVT at term adopt a more differentiated state within the decidua and engage in endocrine and vascular regulation, immune modulation, and stress adaptation. These findings extend the functional role of EVT beyond early pregnancy, suggesting that they contribute to maternal-fetal homeostasis throughout gestation.

Beyond biological insights, our work establishes a gestationally and regionally resolved reference framework for systematic evaluation and reinterpretation of trophoblast organoid models. One key distinction from previous benchmarking studies that focused primarily on villous placenta from a single developmental stage^14,27–29^ is the regional identity of TSC-derived organoids. We show that TSC-Org more closely resemble smooth chorion CTB rather than villous CTB recapitulated in tissue-derived organoids. TSC-Org originate from ITGA6-enriched trophoblast populations^23^, and given the strong expression of *ITGA6* in smooth chorion CTB (Figure S5E), this ITGA6-based enrichment strategy may preferentially select for this population. Recent studies demonstrating derivation of TSC from term smooth chorion further support this interpretation^91^. Together, our findings challenge previous interpretations of TSC-Org as representing villous CTB or EVT progenitor states^27,76^ and instead demonstrate a distinct regional identity, with important implications for their future application. Among tissue-derived models, Term-Org provide a practical alternative given the limited accessibility of early gestational tissues. Although they show weaker *in vivo* trophoblast signatures than FirstTri-Org and only partial acquisition of term-associated gene programs, they broadly recapitulate key villous trophoblast states and therefore remain an informative model. Collectively, these findings highlight the importance of comprehensive *in vivo* reference frameworks for guiding interpretation and application of *in vitro* models.

In summary, our study demonstrates the value of revisiting public single-cell datasets using tailored analytical strategies to uncover new biological insights. By establishing a unified developmental framework of the human placenta in its uterine environment, we provide a foundation for understanding cellular and molecular remodeling across gestation and a resource for investigating pregnancy disorders and improving *in vitro* models of human placentation.

## Acknowledgements

We thank C. Hanna, G.J. Burton, A. Sharkey, and members of the McGovern lab for commenting on the manuscript. This study makes use of data generated by The Chinese University of Hong Kong (CUHK) Circulating Nucleic Acids Research Group, as reported by Tsang et al in Proc Natl Acad Sci USA (DOI: 10.1073/pnas.1710470114). N. McGovern and Q. Li are funded by the Wellcome Trust and Royal Society (G112870) and ERC award funded by UKRI (G122519).

## Author contributions

Q.L., A.M., and N.M. conceived the study. Q.L. developed the computational pipeline, collected data, and performed all analyses. Q.L., A.M., and N.M. wrote the manuscript.

## Methods

### Collection of public scRNA-seq and snRNA-seq datasets of the human placenta

Publicly available scRNA-seq and snRNA-seq datasets of the human placenta and its uterine environment generated using the 10x Genomics platform were collected across three developmental stages: first trimester, second trimester, and term. A total of 10 independent studies were included: Suryawanshi et al.^11^, Arutyunyan et al.^14^, Shannon et al.^13^, Vento-Tormo et al.^10^, Marsh et al.^16^, Tsang et al.^17^, Lu-Culligan et al.^19^, Keenen et al.^28^, Wang et al.^21^, and Yang et al.^18^ (Table S1).

Raw sequencing files (FASTQ or BAM, when FASTQ files were not available) were downloaded from public repositories, including NCBI BioProject, Gene Expression Omnibus (GEO), EMBL-EBI ArrayExpress, and the European Genome-phenome Archive (EGA), under the following accession numbers: PRJNA492324, E-MTAB-12421, GSE174481, E-MTAB-6701, GSE198373, EGAS00001002449, GSE171381, GSE288650, GSE247038, and GSE173193. Associated metadata, including sample identifiers, donor information, gestational age, library preparation protocols, sequencing platforms, and sampling sites, were also collected.

### Preprocessing of collected datasets under a unified framework

To minimize technical variability across studies, all collected scRNA-seq and snRNA-seq datasets were reprocessed from raw sequencing files using a unified analytical framework. A human reference was generated using the GRCh38 genome assembly and gene annotation from GENCODE release 44. Read alignment, gene expression quantification, and cell calling were performed using Cell Ranger (v7.1.0) against the generated reference.

Doublets were initially detected using Scrublet^92^ (v0.2.3), run independently for each sample with default parameters. To further refine doublet identification, two rounds of clustering were performed using 2,000 highly variable genes (HVGs) and 30 principal components (PCs). Clusters showing significantly higher Scrublet doublet scores (Benjamini–Hochberg (BH) adjusted *p*-value < 0.05) were flagged as doublet-enriched.

Ambient RNA contamination was removed using CellBender^93^ (v0.3.0) with a false positive rate of 0.01. A default learning rate of 1×10⁻⁴ was used for most samples; for selected samples requiring improved model convergence, reduced learning rates (5×10⁻⁵, 2.5×10⁻⁵, or 1.25×10⁻⁵) were applied.

To infer cell-of-origin (fetal versus maternal), Souporcell^94^ (v2.5) was run per sample with k = 2 clusters using common variants from the 1000 Genomes Project. Cells and nuclei were excluded from downstream analysis if they met any of the following criteria: (1) ≥15% mitochondrial gene content; (2) <300 genes detected; (3) classified as doublets by Scrublet or Souporcell; (4) removed by CellBender; or (5) unassigned by Souporcell.

### Data integration using a specialised computational pipeline

#### Intra-dataset integration and clustering

Integration across donors was performed separately for each dataset using scVI^95^ (v1.0.3) with 3,000 HVGs and the following parameters: batch_key = "donor", n_layers = 2, n_latent = 20, and gene_likelihood = ’zinb’. Leiden clustering was conducted in the derived latent space using Scanpy^96^ (v1.10.2). Clusters were assigned to broad cell types (trophoblast cells, HBC, fetal fibroblasts, decidual epithelial cells, decidual stromal and perivascular cells, decidual NK and T cells, decidual macrophages, endothelial cells, erythrocytes, and unassigned cells) based on canonical marker gene expression.

To resolve finer subclusters, donor-level integration was repeated separately for each broad cell type using both Harmony^97^ (v1.2.0; 2,000 HVGs and 50 PCs) and scVI (n_layers = 1, n_latent = 10, gene_likelihood = ’zinb’). Leiden clustering was performed on either the 30 Harmony-corrected PCs or the 10 scVI latent dimensions. To maximize detection of rare subtypes, clustering results from Harmony and scVI were combined. Specifically, a Harmony-derived subcluster was further subdivided if >80% of its cells were partitioned into two distinct scVI subclusters, each comprising >20% of cells.

#### Intra-dataset annotation using an automated workflow

To standardize subcluster annotation across datasets, we developed an automated workflow using an iterative merging strategy. Pairwise differential expression analysis was performed across all subclusters. Subclusters lacking significant differentially expressed genes (DEGs) relative to others (BH adjusted *p*-value < 0.05; expression proportion >0.4 in target cluster and <0.1 in comparator cluster) were merged with the most transcriptionally similar subcluster (defined as the one having the fewest genes with BH adjusted *p*-value < 0.05). This process was repeated until no further merges were supported.

Each resulting subcluster was assigned a gene signature defined by genes most distinctively enriched or depleted relative to all other subclusters, considering both differences in expression proportion and recurrence as DEGs across pairwise comparisons. Subclusters whose gene signatures were inconsistent with their assigned broad cell type were excluded as potential doublets.

#### Cross-dataset cell type alignment

To align subclusters across datasets, a similarity matrix was constructed. For each dataset, a CellTypist^98^ (v1.6.3) model was trained based on its annotated subclusters. Each model was then applied to all datasets to obtain the probabilities for each cell being assigned to the subclusters. Similarity between subcluster *i* from study *m* and subcluster *j* from study *n* was quantified using the area under the receiver operating characteristic curve (AUC), where true labels corresponded to cells from study *m* belonging to subcluster *i*, and prediction scores corresponded to probabilities of assignment to subcluster *j* from the model trained on study *n*. Hierarchical clustering of the similarity matrix was used to group transcriptionally concordant subclusters across datasets. Cross-dataset alignment was performed separately for trophoblast cells, stromal/perivascular cells, immune cells, and epithelial/endothelial cells to improve resolution.

#### Cross-dataset integration

To validate cell type alignment across datasets, integration of all datasets was additionally performed (Figure S6A). For trophoblast, stromal/perivascular, and immune cells, scVI was first used to derive a latent representation, followed by a single iteration of Harmony to further mitigate residual batch effects. For epithelial/endothelial cells, Harmony alone was used due to superior integration performance for these cells. Detailed integration parameters for each cell type are provided in Table S2.

#### Final cell annotation

The final set of cell subtypes present across all datasets was determined by combining cross-dataset alignment, integration results, and manual curation. Because certain subclusters were further subdivided during cross-dataset integration, subcluster identities were not directly used as final labels.

For each subtype, high-confidence seed cells were defined based on subclusters showing the strongest transcriptional coherence. Label propagation was then performed in the batch-corrected low-dimensional space. At the broad cell type level, labels were propagated using the ‘LabelPropagation’ function from scikit-learn (v1.5.2), followed by majority voting within subclusters. Additional manual curation was applied for STB, HBC, and erythrocytes to preserve known developmental distinctions.

Subtype-level label propagation was then conducted within each broad cell type using two complementary approaches: (1) semi-supervised propagation using ‘LabelPropagation’, and (2) centroid-based assignment, in which subtype centroids were calculated from seed cells in the integrated low-dimensional space; cells were assigned to the nearest centroid based on Euclidean distance, followed by refinement through majority voting among the 15 nearest neighbors. Both approaches yielded consistent annotations (Figure S6B). The first approach was used for trophoblast cells, and the second for remaining cell types.

### Identification of potential blood contamination

To distinguish tissue-resident immune cells from potential peripheral blood contamination, prior biological knowledge combined with marker gene expression patterns were used to identify blood-derived populations. Fetal neutrophils, which have not been reported to reside in placental tissue, were classified as blood-derived. A distinct NK cell cluster transcriptionally separated from dNK and lacking expression of *NCAM1* was also annotated as blood-derived. Monocyte populations previously characterized as circulating blood monocytes^10^ were similarly annotated as circulating in origin. In addition, pDC and a distinct T cell cluster (bT) expressing high levels of *SELL*, a gene enriched in peripheral blood relative to decidual immune cells in prior single-cell studies, were considered as blood populations. Because B cells of both fetal and maternal origin formed a single transcriptional cluster, tissue-resident and circulating B cells could not be reliably distinguished; therefore, all B cells were conservatively treated as blood-derived.

### Examination of cell abundance changes across gestation

To assess changes in cell abundance across gestation, two complementary approaches were applied. First, for each age point, the proportion of cells belonging to each cell subtype was calculated relative to the total number of cells at that age point. These proportions were then normalized across age points to enable comparison of relative subtype abundance across gestation. Second, differential abundance analysis was performed using the R package miloR^99^ (v2.0.0) at the neighborhood level. Neighborhoods were constructed based on the batch-corrected low-dimensional space derived from scVI and Harmony (see section *Cross-dataset integration*). Differential abundance testing was conducted between first trimester and term samples. Neighborhoods with a spatial false discovery rate (FDR) < 0.1 were considered statistically significant. Neighborhood annotation was performed using the function ‘annotateNhoods’. Neighborhoods in which <40% of cells were assigned to the dominant cell subtype were excluded from subtype-level interpretation.

### Annotation of the stromal and vascular cells in the placenta

Two complementary approaches were used to validate the identity of stromal and vascular cell subtypes identified in the placenta. First, canonical marker genes for fibroblasts, pericytes, SMC, and endothelial subtypes (capillary, arterial, and venous) were curated from the literature, and their expression patterns were examined across placental stromal and vascular populations. Second, a CellTypist model was trained using fibroblast, mural, and endothelial cell annotations from a published organotypic atlas^100^. This trained model was subsequently applied to the placental stromal-vascular compartment to independently predict subtype identities.

### Functional enrichment analysis for fFB subtypes

Marker genes for each fFB subtype were identified using the ‘rank_genes_groups’ function in Scanpy by comparing each subtype to all other cells within the stromal-vascular compartment in the placenta (BH adjusted *p*-value < 0.05, fold change > 2, expression proportion > 0.3). The top 100 marker genes were selected based on the magnitude of differences in expression proportion between the target subtype and all other stromal and vascular cells. Functional enrichment analysis was performed based on the top 100 marker genes for each fFB subtype using Metascape^101^ (v3.5.20260201).

### Prediction of spatial distribution for stromal and vascular cells in the placenta

To infer the spatial localization of fFB, fetal perivascular cells (fPV), and fEC within the mesenchymal core of placental villi, these populations were mapped onto a publicly available spatial transcriptomics dataset profiling early human placenta using STARmap in situ sequencing^15^. Because the spatial data were generated from first-trimester samples, only cells and nuclei derived from first-trimester placental villi were included from the scRNA-seq/snRNA-seq reference, encompassing fFB, fPV, fEC, and other placental cell types (cycCTB, CTB, STB, cycEVT, HBC, and erythrocytes).

Both scRNA-seq/snRNA-seq and spatial data were normalized using SCTransform in Seurat^102^ (v5.1.0). Label transfer from the single-cell/nucleus reference to the query spatial dataset was performed using the anchor-based workflow in Seurat. Briefly, transfer anchors were identified using the ‘FindTransferAnchors’ function based on the top 30 PCs computed from the reference data. Cell type labels were then transferred to the spatial data using the ‘TransferData’ function. Visualization of the predicted identities for one 7 GW sample is shown in Figure 3E.

### Identification of potential cellular interactions in the placenta

Cell-cell communication analysis was performed using CellChat^103^ (v2.2.0) to infer ligand-receptor-mediated interactions among placental cell types (fFB, fPV, fEC, and CTB) based on the curated human ligand-receptor database. Analysis was conducted following the standard CellChat workflow. The RSPO3-LGR5 ligand-receptor pair was manually incorporated due to its established role in trophoblast biology and high expression in the placenta. Selected interactions based on expression level and cell-type specificity are shown in Figure 3F.

### Differential expression analysis between developmental stages for placental stromal and vascular cells

To identify genes showing significant expression changes between first trimester and term in fFB and fPV, differential expression analysis was performed separately for each subtype using the ‘rank_genes_groups’ function in Scanpy. Cells from the first trimester and term samples were compared within each subtype.

Genes meeting the following criteria were considered significantly enriched in either first trimester or term: BH adjusted *p*-value < 0.05, fold change > 2, and expression proportion > 0.3 in the target stage. For each cell subtype, the top 50 first-trimester-enriched genes and top 50 term-enriched genes were selected based on the magnitude of differences in expression proportion between stages. Representative genes reflecting distinct biological processes are shown in Figure 3D.

### Gene set enrichment analysis

To assess enrichment of specific cellular properties within given cell types, associated gene signatures were first identified, and enrichment scores were calculated using the ‘score_genes’ function in Scanpy.

To evaluate proliferative capacity, G2/M and S phase genes were obtained from the cc.genes list in Seurat. To assess similarity to CTB and EVT lineages (Figure 4B), CTB- and EVT-specific gene signatures were defined through differential expression analysis within the trophoblast compartment (CTB, EVT, and STB). For each target cell type, two pairwise comparisons were performed against the other two populations (CTB: CTB vs. STB and CTB vs. EVT; EVT: EVT vs. STB and EVT vs. CTB) to ensure specificity. Genes with BH adjusted *p*-value < 0.01, fold change > 2, and expression proportion > 0.5 in both comparisons were defined as gene signatures. The top 100 genes, ranked by differences in expression proportion between the target cell type and other trophoblast populations, were used as input for ‘score_genes’.

### Trajectory analysis of trophoblast cells

To infer differentiation trajectories across trophoblast populations within the villous placenta, both pseudotime and RNA velocity analyses were performed. Pseudotime inference for either all trophoblast populations or EVT populations alone was conducted using Palantir^104^ (v1.3.3) based on the batch-corrected low-dimensional space (see section *Cross-dataset integration*). The starting cell was defined as the cell showing the highest expression of *LGR5* for analysis of all populations, or *PEG10* for the EVT-restricted analysis. RNA velocity analysis was performed using scVelo^105^ (v0.3.3) in stochastic mode based on 1,000 HVGs after excluding cell cycle-associated genes.

### Spatial mapping of trophoblast cells

To infer the spatial distribution of trophoblast populations (cycCTB, CTB, STB, cycEVT, and EVT), the same publicly available spatial transcriptomics dataset and label transfer strategy described for stromal and vascular cell analysis were applied (see section *Prediction of spatial distribution for stromal and vascular cells in the placenta*). One methodological difference was that both the reference single-cell/single-nucleus data and the query spatial data were normalized using LogNormalize in Seurat, and reciprocal PCA integration was performed on the reference data prior to identification of transfer anchors. Visualization of predicted cell identities for one 9 GW sample is shown in Figure 4D.

To resolve the spatial localization of distinct EVT subtypes at the maternal-fetal interface, a publicly available Visium spatial transcriptomics dataset from early pregnancy was reanalysed^14^. Spatial deconvolution was performed using cell2location^106^ (v0.1.4), with scRNA-seq data from first-trimester villous placenta used as the reference. Reference cell type signatures were estimated using 5,000 HVGs, with parameters batch_key set to “study” and categorical_covariate_keys set to “material” (cell versus nucleus). Parameters for other functions were set according to the recommended workflow in the cell2location tutorial. Estimated cell type abundances per spatial spot were visualized using the ‘plot_spatial’ function in cell2location.

### Functional characterization of EVT subtypes

To characterize functional diversification of EVT subtypes, subtype-specific marker genes, associated gene programs, and predicted interactions with decidual cell types were analyzed.

Marker genes for each EVT subtype were identified using the ‘rank_genes_groups’ function in Scanpy by comparing each subtype with all other EVT. Genes with BH adjusted *p*-value < 0.05, fold change > 2, and expression proportion > 0.3 were considered statistically significant. For each subtype, representative genes reflecting distinct biological processes were selected from the top 100 markers ranked by differences in expression proportion between the target subtype and remaining EVT (Figure 4J).

To infer cellular interactions between EVT and decidual cell populations (dNK, T cells, dM, and EC), ligand-receptor analysis was performed using CellChat as described above (see section *Identification of potential cellular interactions in the placenta*). The IL18-IL18BP ligand-receptor pair was manually incorporated due to the observed upregulation of *IL18BP* in EVT at the late stage of differentiation. Selected interactions corresponding to the biological processes shown in Figure 4J are presented in Figure 4K.

### Collection and reprocessing of public scRNA-seq and snRNA-seq datasets from trophoblast organoid models

Publicly available scRNA-seq and snRNA-seq datasets from distinct trophoblast organoid models (FirstTri-Org, Term-Org, and TSC-Org) generated using the 10x Genomics platform were collected from four studies: Arutyunyan et al.^14^, Shannon et al.^27^, Keenen et al.^28^, and Shannon et al.^13^.

Raw sequencing files (FASTQ, or BAM when FASTQ files were unavailable) were downloaded from EMBL-EBI ArrayExpress and GEO under the following accession numbers: E-MTAB-12650, GSE216244, GSE288650, and GSE174481. Associated metadata, including sample identifiers, donor information, culture medium, library preparation protocols, sequencing platforms, organoid origin (tissue-derived or TSC-derived), and gestational stage of the source tissue (first trimester or term), were also collected.

All raw data were reprocessed under a unified analytical framework as described for the primary tissue (see section *Preprocessing of collected datasets under a unified framework*). Cells and nuclei were excluded from downstream analysis if they met any of the following criteria: (1) ≥15% mitochondrial gene content; (2) <500 genes detected; (3) classified as doublets; (4) removed by CellBender; or (5) annotated as low-quality cells in the original studies.

### Data integration across trophoblast organoid models

To integrate cells and nuclei across organoid models, scVI was applied in combination with Harmony. A latent representation was first learned using scVI, followed by a single iteration of Harmony to further correct residual batch effects between cells and nuclei in the scVI-derived latent space. This procedure was repeated three times, with low-quality cells and nuclei removed at each iteration based on clustering results and manual inspection.

The final uniform manifold approximation and projection (UMAP) visualization shown in Figure 5B was generated using the following parameters: 1,000 HVGs; batch_key = "batch", n_latent = 60, n_layers = 2, max_epochs = 100 in scVI; and batch_key = "material" (cell versus nucleus) in Harmony. Leiden clustering was performed twice. In the first run, clustering was conducted on the Harmony-corrected latent space derived from the parameters described above. In the second run, clustering was performed on a latent space generated under identical settings except that the scVI batch_key was set to "sample". Final Leiden clusters were defined by combining results from both runs: a cluster identified in the first run was subdivided if >80% of its cells were partitioned into two distinct clusters in the second run, each comprising >20% of cells.

### Annotation of Leiden clusters in trophoblast organoids

Gene signatures for CTB, STB, EVT, and scCTB, derived from primary tissue, were used to annotate Leiden clusters identified in the organoid datasets. For each cell type, pairwise differential expression analysis was performed on primary tissue data by comparing the target cell type with each of the remaining trophoblast populations using the ‘rank_genes_groups’ function in Scanpy. Genes meeting the following criteria in all pairwise comparisons were defined as cell type-specific signatures: BH adjusted *p*-value < 0.05, fold change > 2, and expression proportion > 0.3. The top 50 signature genes were then selected based on the magnitude of difference in expression proportion between the target cell type and all other trophoblast populations. These gene sets from each cell type were used to infer cell identities in the organoids by computing per-cell scores using the ‘score_genes’ function in Scanpy.

### Cell type alignment between organoids and primary tissue

To further validate cell identities in the organoids, two complementary label transfer approaches were applied using primary tissue as the reference. First, a CellTypist model was trained on all annotated trophoblast subtypes from primary tissue and subsequently used to predict cell identities in the organoids. Second, scANVI^107^ was applied to leverage primary tissue annotations for semi-supervised label transfer. A scVI model was first trained on the combined dataset (primary tissue and organoids) using the union of HVGs from both sources, with the following parameters: batch_key = "donor", n_layers = 2, n_latent = 60, and max_epochs = 300. A scANVI model was next initialized using weights from the scVI model and further trained with max_epochs = 25 and n_samples_per_label = 300. The resulting scANVI model was subsequently used to predict cell identities in the organoids.

### Assessment of developmental stage identity in tissue-derived organoids

To determine which developmental stage tissue-derived organoids (FirstTri-Org and Term-Org) most closely resemble, stage-specific gene signatures were first defined for CTB and STB separately by comparing first-trimester and term cells/nuclei from primary tissue.

For STB, only single nuclei were included in cross-stage comparisons. Stage-enriched genes (first trimester-enriched or term-enriched) were defined as those meeting the following criteria: BH adjusted *p*-value < 0.01, fold change > 2, and expression proportion > 0.5 in the target stage. For each stage, the top 50 enriched protein-coding genes ranked by differences in expression proportion between stages were selected and used as input for the ‘score_genes’ function in Scanpy to evaluate STB populations in organoids.

For CTB, only cells were included in differential expression analysis to minimize potential confounding effects from input type (cell versus nucleus). Stage-enriched genes were defined using the following criteria: BH adjusted *p*-value < 0.01, fold change > 1.75, expression proportion > 0.3 in the target stage, and < 0.15 in the comparator stage. Up to 50 top enriched protein-coding genes per stage were selected based on differences in expression proportion between stages. To assess how organoids align with *in vivo* developmental stages, the expression patterns of first trimester-enriched and term-enriched CTB genes were then examined in FirstTri-Org and Term-Org. For each gene set, the proportion of genes expressed in CTB from each organoid model (expression proportion > 0.3) was calculated. Differential expression analysis between CTB from FirstTri-Org and Term-Org was additionally performed, and genes were considered significantly different between models if they met the following criteria: BH adjusted *p*-value < 0.01, fold change > 1.75, and expression proportion > 0.3 in the target model.

---

**Figure S1.**
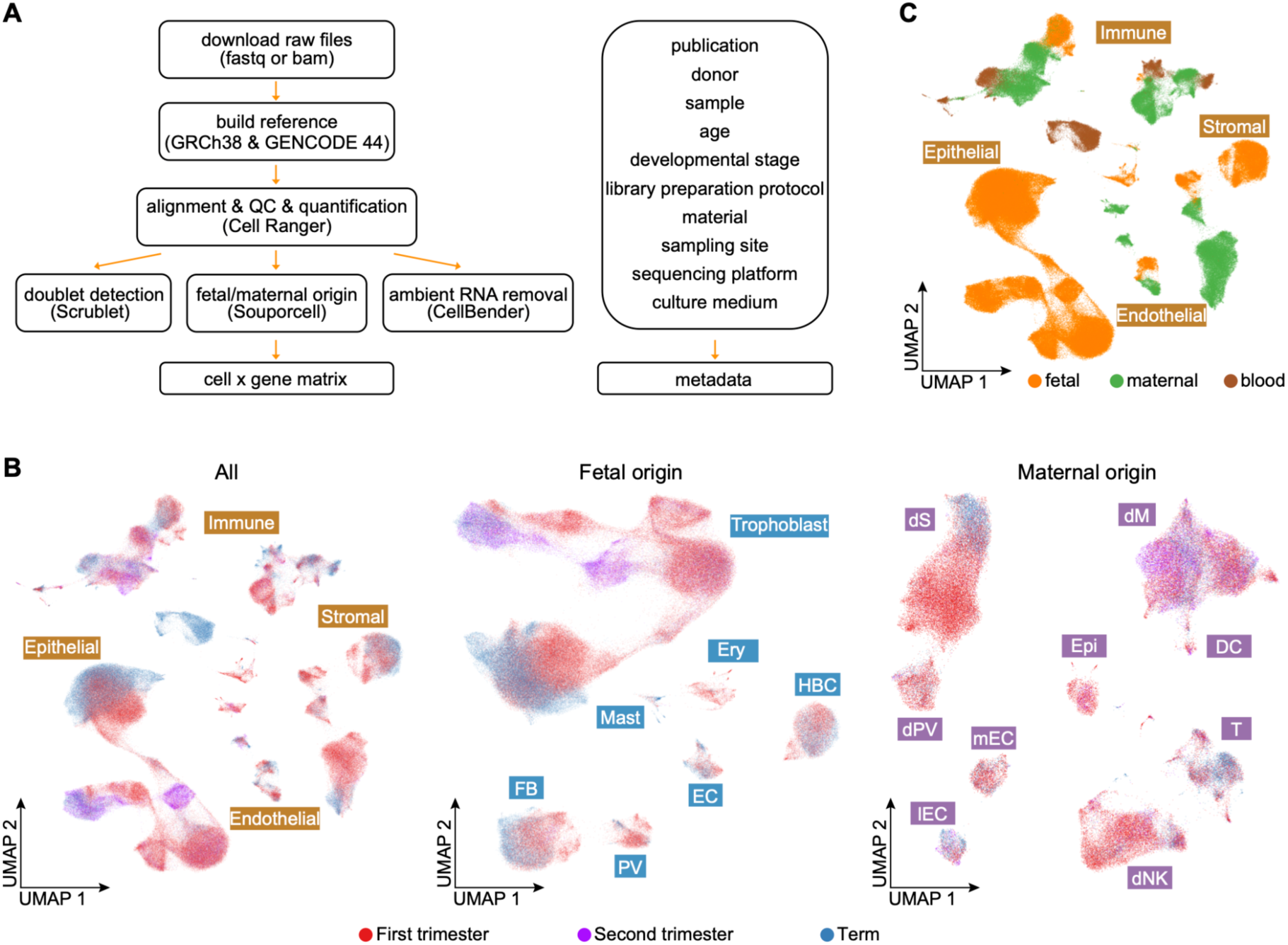
A unified preprocessing framework and quality control of all datasets collected. (**A**) A unified preprocessing framework used to reprocess all scRNA-seq and snRNA-seq datasets from raw data, including cell filtering, doublet identification, ambient RNA removal, maternal-fetal origin inference, as well as metadata collection. (**B**) UMAP visualization of all cells and nuclei (left), as well as fetal-origin (middle) and maternal-origin (right) populations, colored by gestational stages. (**C**) UMAP visualization of all cells and nuclei colored by cell origin.

**Figure S2.**
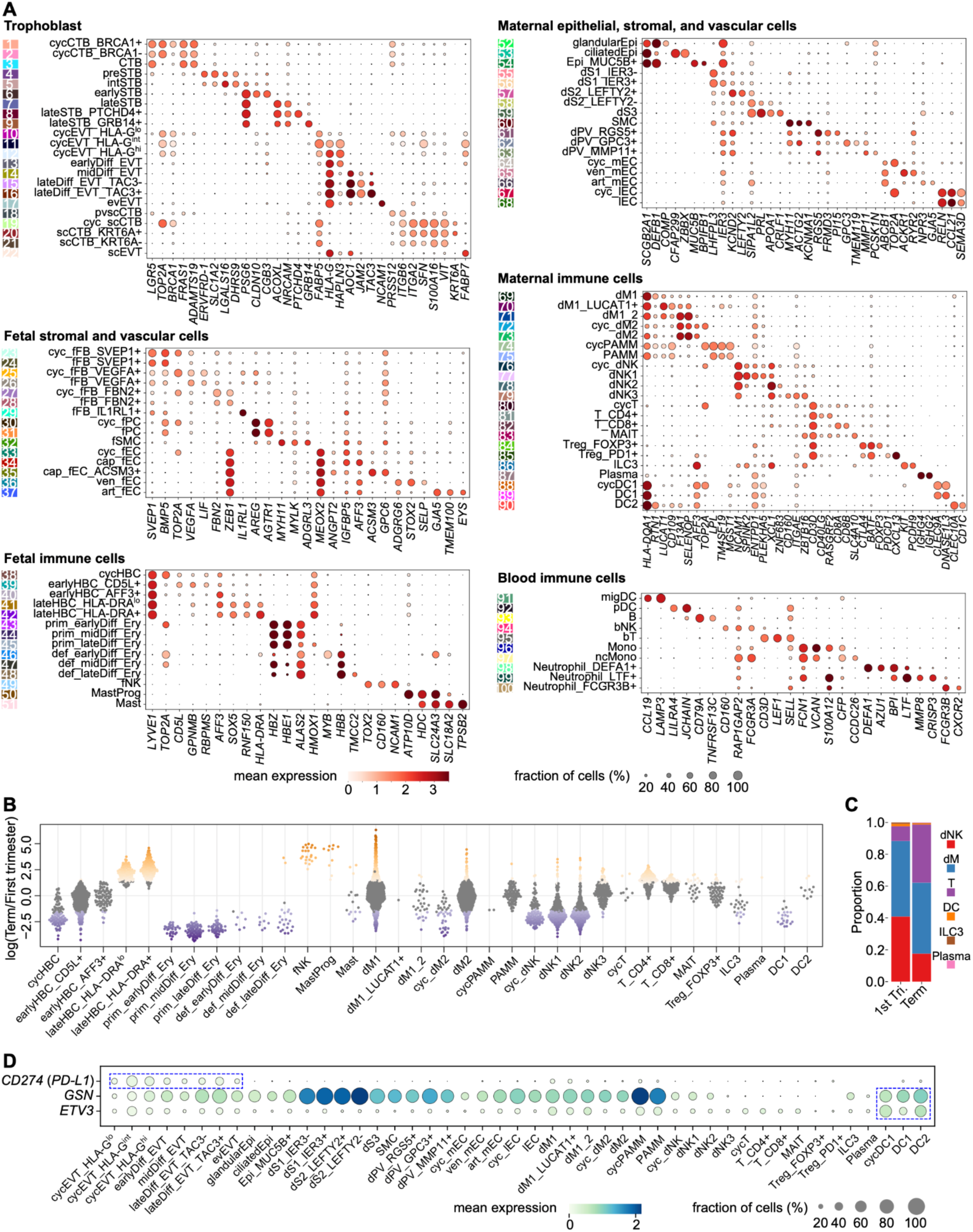
Cell subtype gene signatures and abundance changes across gestation. (**A**) Dot plot showing expression of gene signatures defining the 100 cell subtypes identified in the human placenta and its uterine environment across gestation. Numbers on the left correspond to those shown in Figure 2A. Dot color and size indicate average expression level and proportion of cells expressing the genes, respectively. (**B**) Changes in abundance of fetal and maternal immune subtypes between the first trimester and term. Each dot represents a neighborhood assigned to a subtype, and colored dots indicate significant abundance changes (FDR < 0.1). Only subtypes with at least one neighborhood assigned are shown. (**C**) Proportions of individual immune populations among decidual immune cells in the first trimester and term. (**D**) Dot plot showing expression of *CD274* (*PD-L1*), *GSN* (Gelsolin), and *ETV3* across cell subtypes in the cell column and decidua. Dot color and size indicate average expression level and proportion of cells expressing the genes, respectively. Dotted blue rectangles highlight expression of *CD274* in EVT and *GSN* and *ETV3* in DC populations.

**Figure S3.**
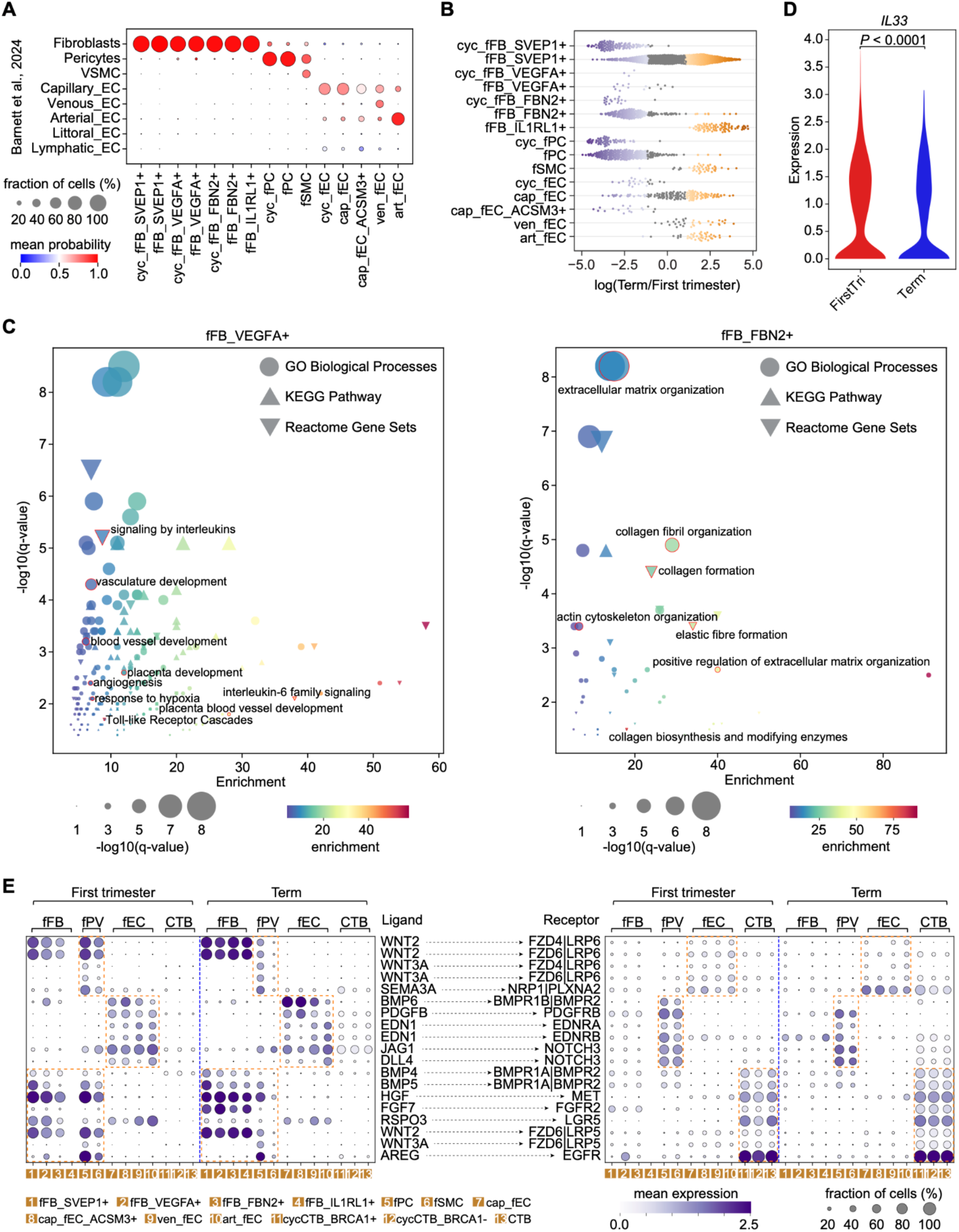
Placental stromal and vascular cells show dynamic remodelling across gestation. (**A**) Predicted cell identities for placental stromal and vascular subtypes based on a public single-cell atlas from Barnett et al., 2024. Dot color and size indicate prediction probability and proportion of cells from this study aligned to each reference subtype, respectively. (**B**) Changes in abundance of placental stromal and vascular subtypes between the first trimester and term. Each dot represents a neighborhood assigned to a subtype, and colored dots indicate significant abundance changes (FDR < 0.1). (**C**) GO biological processes, KEGG and Reactome pathways enriched in the top 100 marker genes for *VEGFA*+ fFB (left) and *FBN2*+ fFB (right) subtypes. Vascular- and ECM-associated terms are highlighted. Color gradient and dot size indicate enrichment scores and BH adjusted *p*-values, respectively. (**D**) Violin plot showing expression of *IL33* in fetal endothelial cells from first trimester and term samples. *P*-value calculated between the two stages by a *t* test is shown on top. (**E**) Ligand-receptor-mediated interactions between stromal, vascular, and trophoblast populations in the placenta, as shown in Figure 3F. Expression of the ligands (left) and receptors (right) across cell subtypes is separated by gestational stages (first trimester versus term). Dot color and size indicate average expression level and proportion of cells expressing the genes, respectively.

**Figure S4.**
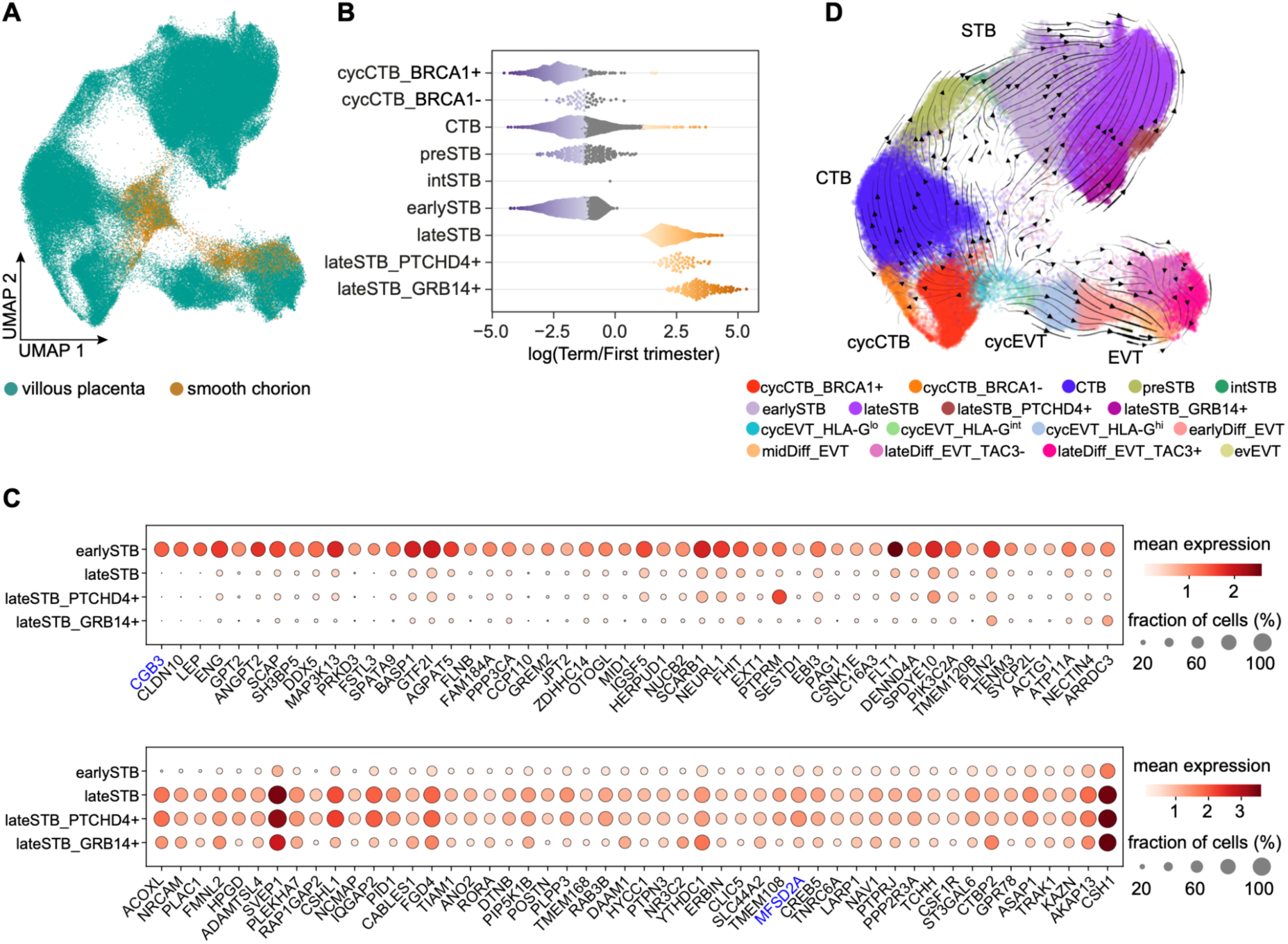
Dynamic remodeling of the trophoblast lineage across gestation. (**A**) UMAP visualization of trophoblast cells colored by placental regions. (**B**) Changes in abundance of CTB and STB subtypes between the first trimester and term. Each dot represents a neighborhood assigned to a subtype, and colored dots indicate significant abundance changes (FDR < 0.1). (**C**) Dot plot showing expression of the top 50 protein-coding genes enriched in first-trimester STB (top) and term STB (bottom). The two genes associated with hCG production and DHA transport are highlighted in blue. Dot color and size indicate average expression level and proportion of cells expressing the genes, respectively. (**D**) RNA velocity analysis showing the inferred differentiation trajectory of villous trophoblast cells.

**Figure S5.**
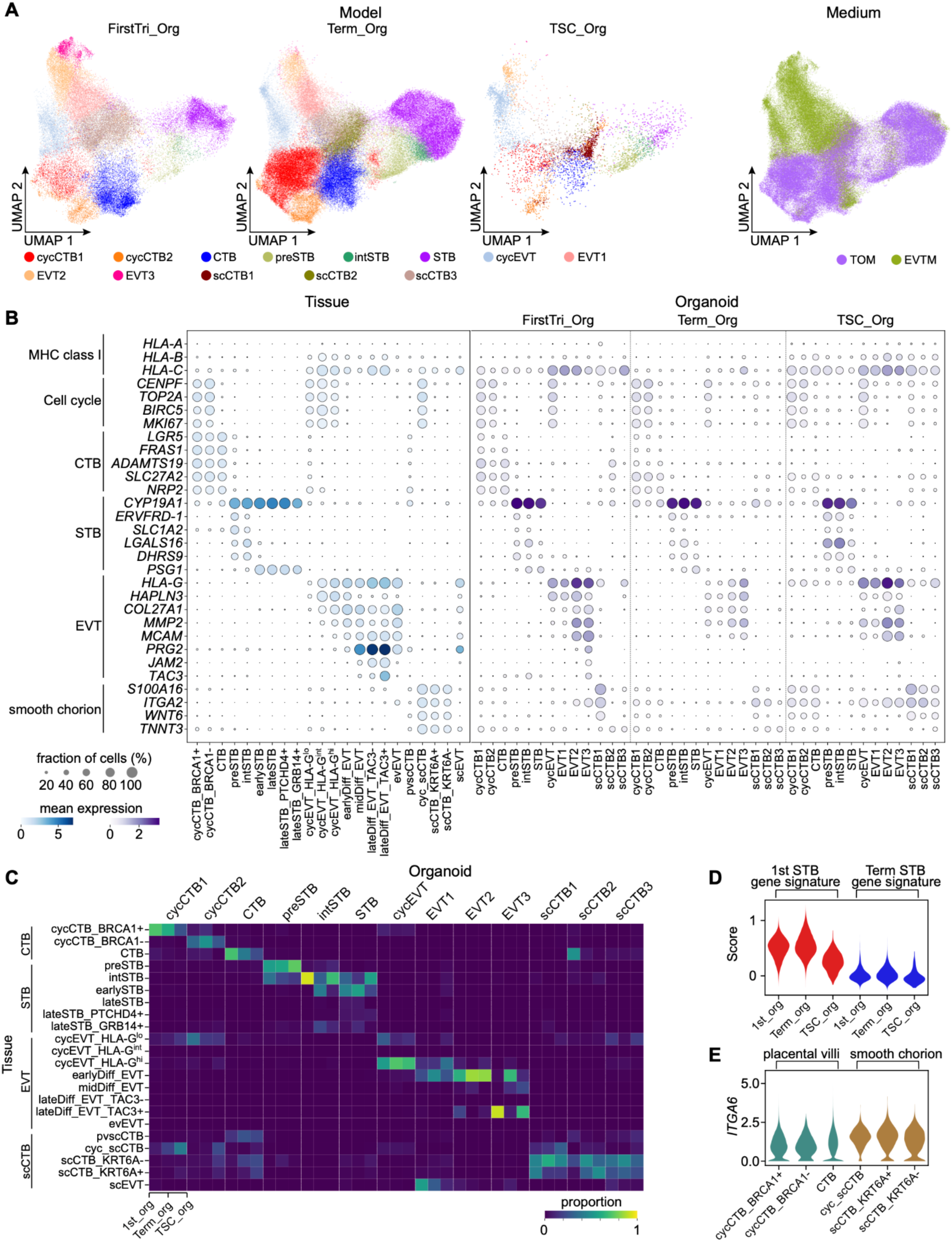
Systematic benchmarking of trophoblast organoid models. (**A**) UMAP visualization of trophoblast subtypes identified in the three organoid models (FirstTri-Org, Term-Org, and TSC-Org), with cells colored by cell subtypes and culture media, respectively. TOM, trophoblast organoid medium; EVTM, EVT differentiation medium. (**B**) Dot plot showing expression of MHC class I genes, cell cycle-associated genes, and *in vivo* lineage gene signatures across trophoblast subtypes from primary tissue and the three organoid models. Dot color and size indicate average expression level and proportion of cells expressing the genes, respectively. (**C**) Heatmap showing alignment of cell subtypes between primary tissue and the three organoid models. Color gradient indicates the proportion of cells from each organoid model aligned to each *in vivo* subtype from scANVI. (**D**) Violin plot showing distributions of first-trimester STB and term STB gene signature scores in *in vitro* STB across organoid models. Scores were calculated based on expression of the top 50 protein-coding genes enriched in first-trimester or term STB identified *in vivo*. (**E**) Violin plot showing expression of *ITGA6* across villous CTB and smooth chorion CTB subtypes.

**Figure S6.**
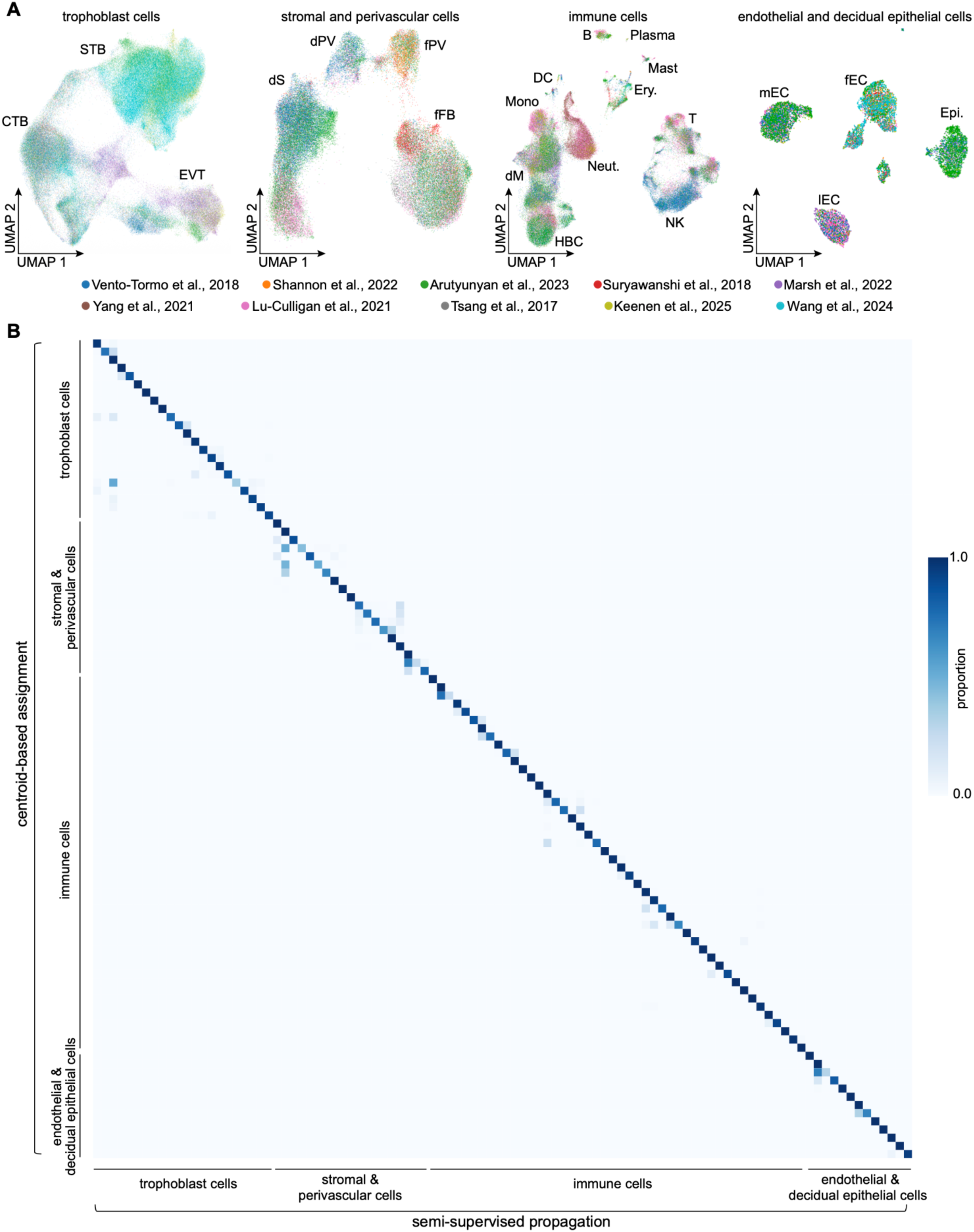
Cross-dataset alignment and integration for final cell annotation. (**A**) UMAP visualization of trophoblast, stromal/perivascular, immune, and endothelial/epithelial populations, showing cross-dataset integration within each cellular compartment. Cells and nuclei are colored by datasets. Neut., neutrophil. (**B**) Heatmap showing highly concordant cell annotations generated by two complementary label propagation approaches: centroid-based assignment (y axis) and semi-supervised propagation (x axis). Color gradient indicates the proportion of cells/nuclei shared between the two approaches for each cell subtype.

**Table S1.** Metadata for public scRNA-seq and snRNA-seq datasets collected.

| Study | Donor | Stage | Age (GW) | Region | Material | Accession no. |
| --- | --- | --- | --- | --- | --- | --- |
| Suryawanshi et al., 2018 | Pla17 | 1<sup>st</sup> tri. | 6W4D | villi | scRNA | PRJNA492324 |
|  | Pla23 | 1<sup>st</sup> tri. | 7W6D | villi, decidua | scRNA |  |
| Arutyunyan et al., 2023 | Hrv100 | 1<sup>st</sup> tri. | 7W | villi, decidua | scRNA | E-MTAB-12421 |
|  | Hrv99 | 1<sup>st</sup> tri. | 10W | villi, decidua | scRNA |  |
|  | P13 | 1<sup>st</sup> tri. | 10-11W | villi, decidua | snRNA |  |
|  | Hrv98 | 1<sup>st</sup> tri. | 11W | villi, decidua | scRNA |  |
| Shannon et al., 2022 | P244 | 1<sup>st</sup> tri. | 6.2W | villi | scRNA | GSE174481 |
|  | P89 | 1<sup>st</sup> tri. | 6.4W | villi | scRNA |  |
|  | P277 | 1<sup>st</sup> tri. | 6.4W | villi | scRNA |  |
|  | P295 | 1<sup>st</sup> tri. | 6.5W | villi | scRNA |  |
|  | P21 | 1<sup>st</sup> tri. | 11.2W | villi | scRNA |  |
|  | P130 | 1<sup>st</sup> tri. | 11.4W | villi | scRNA |  |
|  | P899 | 1<sup>st</sup> tri. | 11.4W | villi | scRNA |  |
| Vento-Tormo et al., 2018 | D11 | 1<sup>st</sup> tri. | 6-7W | villi | scRNA | E-MTAB-6701 |
|  | D8 | 1<sup>st</sup> tri. | 8W | villi, decidua | scRNA |  |
|  | D10 | 1<sup>st</sup> tri. | 9-10W | villi, decidua | scRNA |  |
|  | D12 | 1<sup>st</sup> tri. | 10W | villi, decidua | scRNA |  |
|  | D7 | 1<sup>st</sup> tri. | 10W | decidua | scRNA |  |
|  | D6 | 1<sup>st</sup> tri. | 12W | decidua | scRNA |  |
|  | D9 | 1<sup>st</sup> tri. | 12W | villi, decidua | scRNA |  |
| Marsh et al., 2022 | SP1 | 2<sup>nd</sup> tri. | 17W6D | villi | scRNA | GSE198373 |
|  | SP2 | 2<sup>nd</sup> tri. | 18W2D | smooth chorion | scRNA |  |
|  | SP3 | 2<sup>nd</sup> tri. | 23W | villi, smooth chorion | scRNA |  |
|  | SP4 | 2<sup>nd</sup> tri. | 24W | villi, smooth chorion | scRNA |  |
| Tsang et al., 2017 | M12475 | term | 38W | villi | scRNA | EGAS00001002449 |
|  | M12551 | term | 38W | villi | scRNA |  |
|  | M12491 | term | 38W | villi | scRNA |  |
|  | M12548 | term | 38W2D | villi | scRNA |  |
| Lu-Culligan et al., 2021 | C3 | term | 39W | decidua | scRNA | GSE171381 |
|  | C2 | term | 39W | villi | scRNA |  |
|  | C1 | term | 39W | villi | scRNA |  |
| Keenen et al., 2025 | AGPL4 | term | 37W | villi, decidua | sc/snRNA | GSE288650 |
|  | AGPL5 | term | 38W | villi, decidua | snRNA |  |
|  | AGPL3 | term | 39W | villi, decidua | sc/snRNA |  |
|  | AGPL2 | term | 39W | villi, decidua | scRNA |  |
| Wang et al., 2024 | donor4 | 1<sup>st</sup> tri. | 6W | villi | snRNA | GSE247038 |
|  | donor3 | 1<sup>st</sup> tri. | 7W | villi | snRNA |  |
|  | donor5 | 1<sup>st</sup> tri. | 8W | villi | snRNA |  |
|  | donor1 | 1<sup>st</sup> tri. | 8W | villi | snRNA |  |
|  | donor2 | 1<sup>st</sup> tri. | 8W | villi | snRNA |  |
|  | donor6 | 1<sup>st</sup> tri. | 9W | villi | snRNA |  |
|  | donor12 | term | 38W | villi | snRNA |  |
|  | donor10 | term | 38W | villi | snRNA |  |
|  | donor9 | term | 38W | villi | snRNA |  |
|  | donor8 | term | 38W | villi | snRNA |  |
|  | donor11 | term | 38W | villi | snRNA |  |
|  | donor7 | term | 39W | villi | snRNA |  |
| Yang et al., 2021 | YC2 | term | 38W3D | villi | scRNA | GSE173193 |
|  | YC1 | term | 40W4D | villi | scRNA |  |

**Table S2.** Detailed parameters for cross-dataset integration.

| Cell type | Figure | HVG selection |  |  | scVI |  |  | Harmony |  |
| --- | --- | --- | --- | --- | --- | --- | --- | --- | --- |
| | | #HVGs | Batch key | rmCC | Batch key | $n_{\text{latent}}$ | $n_{\text{epoch}}$ | Batch key | $n_{\text{iter}}$ |
| All | Figure 2A | 3000 | stage | N | donor | 60 | 100 | study | 1 |
| Fetal | Figure 2B | 2000 | donor | N | donor | 60 | 100 | study | 1 |
| Maternal | Figure 2C | 3000 | donor | N | donor | 60 | 100 | study | 1 |
| Stromal & vascular | Figure 3A | 3000 | stage | Y | donor | 60 | 100 | study | 1 |
| Trophoblast | Figures 4A & S6A | 2000 | study | N | donor | 60 | 100 | study | 1 |
| Organoids | Figure 5B | 1000 | sample | N | batch | 60 | 100 | material | 1 |
| Stromal/PV | Figure S6A | 2000 | study | Y | donor | 30 | 100 | study | 1 |
| Immune | Figure S6A | 2000 | stage | N | donor | 60 | 100 | study | 1 |
| EC/Epi | Figure S6A | 1000 | stage | N | \ |  |  | study & donor | Conv. |

HVG: highly variable gene rmCC: remove cell cycle-associated genes Y: yes N: no $n_{\text{latent}}$ : dimensionality of the latent space $n_{\text{epoch}}$ : number of epochs $n_{\text{iter}}$ : number of iterations PV: perivascular cells EC: endothelial cells Epi: epithelial cells Conv.: converge

